# Mapping threatened Thai bovids provides opportunities for improved conservation outcomes in Asia

**DOI:** 10.1101/2023.08.25.554763

**Authors:** Wantida Horpiencharoen, Renata L. Muylaert, Jonathan C. Marshall, Reju Sam John, Antony J. Lynam, Alex Riggio, Alexander Godfrey, Dusit Ngoprasert, George A. Gale, Eric Ash, Francesco Bisi, Giacomo Cremonesi, Gopalasamy Reuben Clements, Marnoch Yindee, Nay Myo Shwe, Chanratana Pin, Thomas N. E. Gray, Saw Soe Aung, Seree Nakbun, Stephanie G. Manka, Robert Steinmetz, Rungnapa Phoonjampa, Naret Seuaturien, Worrapan Phumanee, David T. S. Hayman

## Abstract

Wild bovids provide important ecosystem functions throughout their ranges. Five wild bovids remain in Thailand: gaur (*Bos gaurus*), banteng (*Bos javanicus*), wild water buffalo (Bubalus arnee), mainland serow (*Capricornis sumatraensis*) and Chinese goral (*Naemorhedus griseus*). However, their populations and habitats have declined substantially and become fragmented. Here, we identify potentially suitable habitat for these threatened bovids using ecological niche models and quantify how much suitable area remains within protected areas. We combined species occurrence data with environmental variables and used spatially-restricted Biotic-Abiotic-Mobility frameworks with species-specific and single large accessible areas. We used ensembles from eight algorithms for generating maps and out-of-sample predictions to validate model performance against new data. Gaur, banteng, and buffalo models performed well throughout the entire distribution (≥62%) and in Thailand (≥80%). Mainland serow and Chinese goral performed poorly for the entire distribution and in Thailand, though a 5 km movement buffer markedly improved model performance for serow. Particularly large suitable areas were in Thailand and India for gaur, Cambodia and Thailand for banteng, and India for buffalo. Over 50% of overall suitable habitat is located outside protected areas, with just 9% for buffalo in Thai protected areas, highlighting area for potential habitat management and conflict mitigation.

## Introduction

An important task of wildlife research and conservation is to define the distributional ecology of species and to understand how they relate to the environment, climate and other organisms (Franklin, 2009). Ecological niche models (ENM) are applied to predict the geographic distribution suitable for a species by using ecological niche dimensions combined with species’ presence data (Soberon & Peterson, 2005). ENM can be approached using the ‘Biotic-Abiotic-Mobility’ (BAM) framework, which considers the relationship between the species’ distribution, geographical and climatic factors and explains the influence of factors on predicted habitat suitability (Peterson & Soberón, 2012). Abiotic (A) factors generally determine the potential distribution (or fundamental niche) of a species, and the intersection of abiotic and biotic (B) factors form the realised niche, or the part of this potential distribution where species actually live (Soberón & Nakamura, 2009). Mobility (M) is the area accessible by species related to their distribution over periods of time (the ‘accessible area’; (Barve et al., 2011)). Selecting the extent of species’ accessible areas, including buffer zones, impacts model prediction results (Anderson & Raza, 2010; Barve et al., 2011).

Wild Bovidae (Mammalia: Artiodactyla) play significant ecological roles in tropical forests and grasslands (Hassanin, 2014). Bovids are grazers and browsers, modifying plant diversity and abundance within ecosystems (Ripple et al., 2015; Romero et al., 2015). Large wild bovids are also the prey of predators such as tigers (*Panthera tigris*) and leopards (*Panthera pardus*) (Simcharoen et al., 2018). Throughout Asia, wild bovid populations are threatened by poaching (Gray et al., 2018) and habitat loss (Nguyen, 2009), especially in South to Southeast Asia (Giam & Wilcove, 2012). Natural habitats have been disturbed by free-grazing livestock, which can lead to interbreeding (e.g. between domestic and wild water buffalo, (Kaul et al., 2019), increased competition for food and natural resources (Bhandari et al., 2022), and increased risk of disease transmission between wildlife and livestock (Hassell et al., 2017). Moreover, habitat destruction is likely to influence the species’ distribution and behaviour adaptation, which could lead to shared natural resources and conflict between humans and wild bovids.

In South and Southeast Asia, there are 27 recognised bovid species (IUCN, 2021), of which seven species are listed as vulnerable, five as endangered and three as critically endangered with extinction. Thailand has five bovid species (gaur; *Bos gaurus*, banteng; *Bos javanicus*, wild water buffalo; *Bubalus arnee*, mainland serow; *Capricornis sumatraensis* and Chinese goral; *Naemorhedus griseus*) remaining in their natural habitat. These species are distributed in other countries from South to Southeast Asia (Figure 1) and also have different suitable habitats. For example, gaur can be found in evergreen forest or grassland and range from India, Nepal, across Southeast Asia to Peninsula Malaysia (Duckworth et al., 2016). Mainland serow also has a wide distribution from Nepal to Sumatra in Indonesia through hill forests to shrubland habitats (Phan et al., 2020). Nevertheless, the prediction of the remaining habitat quality and suitability in Thailand and other countries have been conducted only in some protected areas (Chaiyarat et al., 2019; Pintana & Lakamavichian, 2013), but not at the regional or national level.

**Figure 1.**
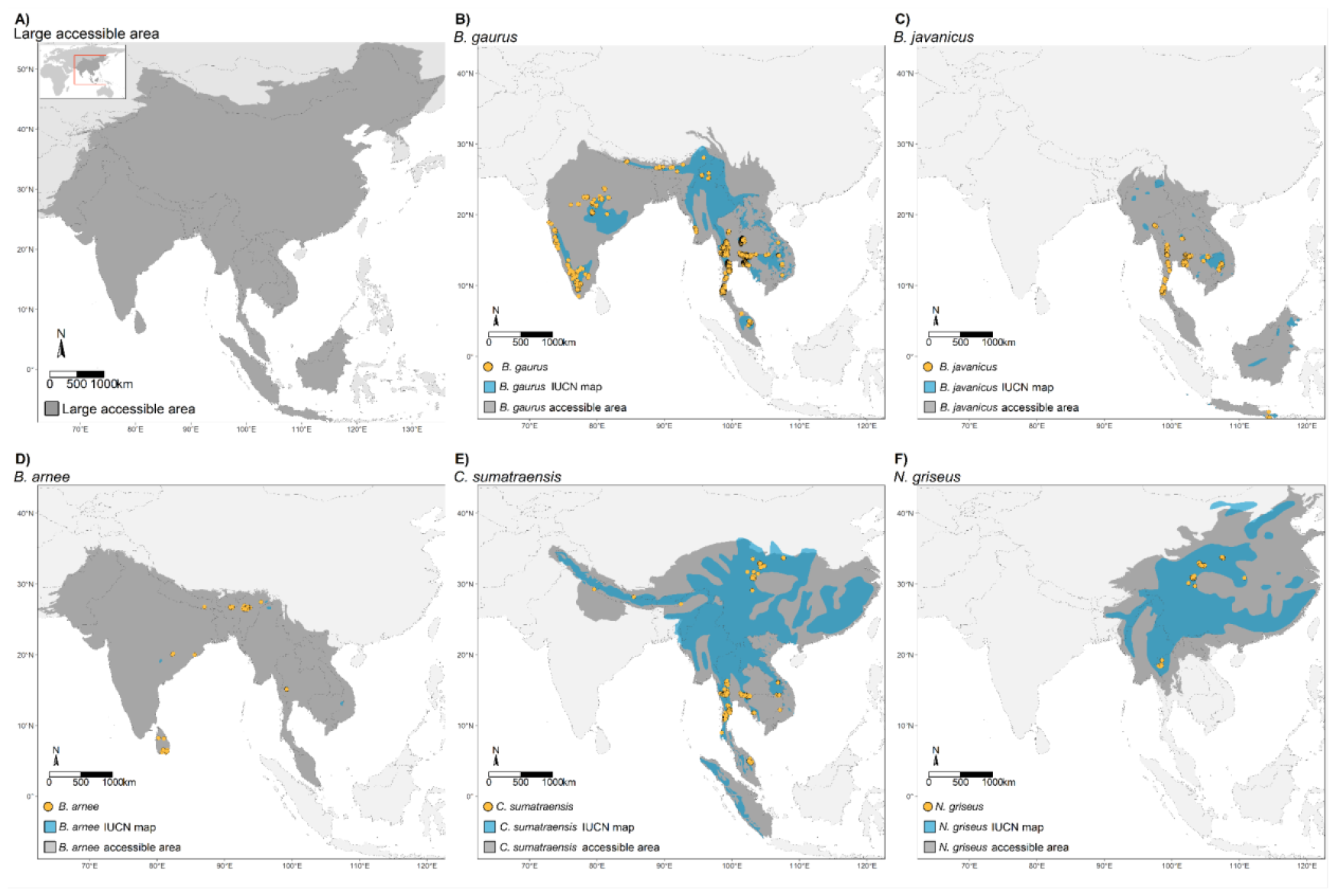
Species occurrence data before thinning (yellow circles), IUCN polygons (blue areas) and study areas (grey areas) used in model building for five wild bovid species. First, a common large ‘accessible area’ (A) was used for all species for model building, and then species-specific accessible areas (B-F) for individual species.**Species Occurrence data**

Species distribution modelling provides an overview of potential habitats for threatened species and aids in conservation planning (Catullo et al., 2008). For instance, previous studies have focused on identifying potentially high-quality habitat connectivity and fragmentation (Crooks et al., 2011) as well as predicting global biodiversity trends (Araújo et al., 2019). In Thailand, there are several studies that have predicted habitat suitability for some of these five wild bovids in local areas (Prayoon et al., 2021), but habitat suitability studies for larger extents across their distribution are lacking.

Here, we built ENM for the five Thai wild bovid species: gaur, banteng, wild water buffalo, mainland serow and Chinese goral at two scales: first, at the regional scale throughout the entire distribution and, second, at the country scale in Thailand. We aim to 1) identify the potential distribution for these five species in South to Southeast Asia, and 2) identify conservation areas in their geographical distribution, with a particular focus on Thailand.

## Materials and methods

Our workflow consisted of two main processes of data preparation and model building (summarised in Figure S1) that generated habitat suitability maps for all species and accessible areas used. Data preparation consisted of gathering the species occurrence data and environmental data and selecting the accessible areas. Then, the model building consisted of pre-processing, processing and post-processing steps.

### Study area

The study area consists of 13 Asian countries: Bhutan, Bangladesh, Cambodia, China, India, Indonesia, Laos, Malaysia, Myanmar, Nepal, Sri Lanka, Thailand and Vietnam (Figure 1), that cover the distribution of gaur, banteng, wild water buffalo, mainland serow and Chinese goral based on the literature (Table S1).

We compiled species occurrence data collected from GPS records collected between January 2000 and June 2021 from researchers, government, NGOs (World Wildlife Fund, Wildlife Conservation Society, Freeland [Ash et al. (2021)], Panthera, Fauna & Flora International, Friends of Wildlife and RIMBA) and open data sources, including GBIF (https://www.gbif.org/) and eMammal (https://emammal.si.edu/). The data coverage by country can be found in Table 1. The occurrence data were collected through observation of animal signs (e.g. footprint and dung) during forest patrols, direct observation during wildlife surveys, camera trapping, and radio-collar signals (Table S2). We used only the research grade observation for GBIF data, which included the photo for species identification. We filtered all the occurrences and excluded occurrence records outside the species-specific accessible area, duplicated records from the same species and museum collections.

**Table 1.**
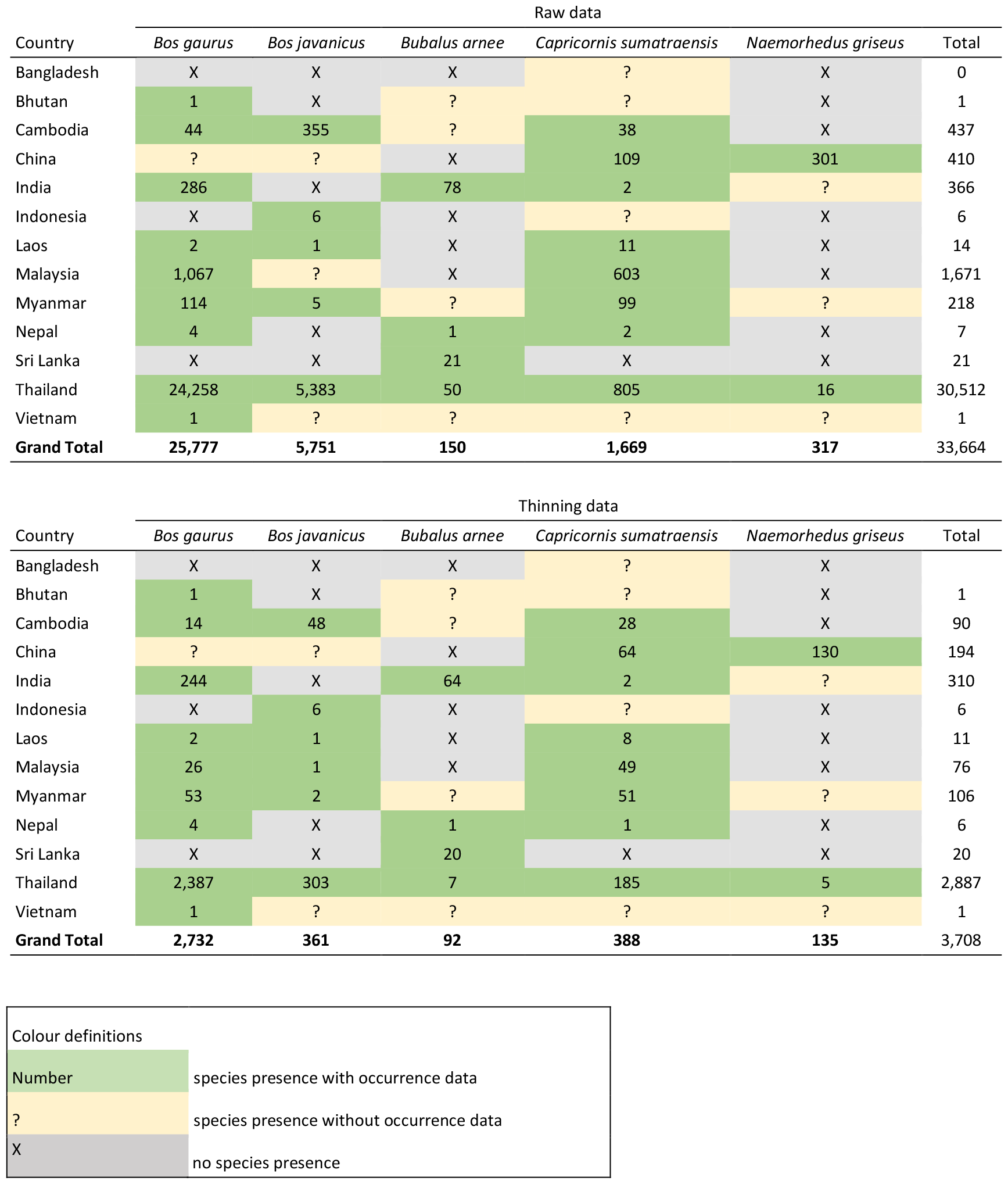
The number of raw and after spatial thinning occurrence points is shown by species and country.

| Raw data |  |  |  |  |  |  |
| --- | --- | --- | --- | --- | --- | --- |
| Country | *Bos gaurus* | *Bos javanicus* | *Bubalus arnee* | *Capricornis sumatraensis* | *Naemorhedus griseus* | Total |
| Bangladesh | X | X | X | ? | X | 0 |
| Bhutan | 1 | X | ? | ? | X | 1 |
| Cambodia | 44 | 355 | ? | 38 | X | 437 |
| China | ? | ? | X | 109 | 301 | 410 |
| India | 286 | X | 78 | 2 | ? | 366 |
| Indonesia | X | 6 | X | ? | X | 6 |
| Laos | 2 | 1 | X | 11 | X | 14 |
| Malaysia | 1,067 | ? | X | 603 | X | 1,671 |
| Myanmar | 114 | 5 | ? | 99 | ? | 218 |
| Nepal | 4 | X | 1 | 2 | X | 7 |
| Sri Lanka | X | X | 21 | X | X | 21 |
| Thailand | 24,258 | 5,383 | 50 | 805 | 16 | 30,512 |
| Vietnam | 1 | ? | ? | ? | ? | 1 |
| **Grand Total** | **25,777** | **5,751** | **150** | **1,669** | **317** | **33,664** |

| Thinning data |  |  |  |  |  |  |
| --- | --- | --- | --- | --- | --- | --- |
| Country | *Bos gaurus* | *Bos javanicus* | *Bubalus arnee* | *Capricornis sumatraensis* | *Naemorhedus griseus* | Total |
| Bangladesh | X | X | X | ? | X |  |
| Bhutan | 1 | X | ? | ? | X | 1 |
| Cambodia | 14 | 48 | ? | 28 | X | 90 |
| China | ? | ? | X | 64 | 130 | 194 |
| India | 244 | X | 64 | 2 | ? | 310 |
| Indonesia | X | 6 | X | ? | X | 6 |
| Laos | 2 | 1 | X | 8 | X | 11 |
| Malaysia | 26 | 1 | X | 49 | X | 76 |
| Myanmar | 53 | 2 | ? | 51 | ? | 106 |
| Nepal | 4 | X | 1 | 1 | X | 6 |
| Sri Lanka | X | X | 20 | X | X | 20 |
| Thailand | 2,387 | 303 | 7 | 185 | 5 | 2,887 |
| Vietnam | 1 | ? | ? | ? | ? | 1 |
| **Grand Total** | **2,732** | **361** | **92** | **388** | **135** | **3,708** |

| Colour definitions |  |
| --- | --- |
| Number | species presence with occurrence data |
| ? | species presence without occurrence data |
| X | no species presence |

### Environmental variables

Hypothesized environmental variables were selected based on species’ habitat and distribution related literature Table S1. We used 28 variables (supplementary material, Table S3) for model construction, including 19 bioclimatic variables (Booth et al., 2014) (average for 1970-2000) from WorldClim v2 (Fick & Hijmans, 2017), elevation (Shuttle Radar Topography Mission-SRTM) from WorldClim (Fick & Hijmans, 2017), slope (Amatulli et al., 2020), five land cover fractions (grass, tree, urban, water and crop) (Buchhorn et al., 2019), human population density (Stevens et al., 2015) and greenness through the normalized difference vegetation index (NDVI) (Didan, 2015). All layers were processed using the geographic coordinates system (Datum WGS84) and ∼1 km^2^ spatial resolution. We transformed the human population density using logarithm base 10 to adjust for skewness. We rescaled the NDVI layer by multiplying all values with a scale factor (0.0001), based on the Moderate Resolution Imaging Spectroradiometer (MODIS) User’s guidelines (Didan et al., 2015).

### Accessible areas

The accessible area refers to the parts of the world accessible to species via dispersal over time (Barve et al., 2011). The extent of the accessible area and the inclusion of a buffer zone have an important effect on ENM performance (Anderson & Raza, 2010; Barve et al., 2011). We used two accessible area sizes to delimitate our modelling extent (Figure 1). The first larger accessible area (hereon LA) includes most of the Asian continent and its ecoregions, and all species distributions are included as a common extent. The second accessible area was more restricted and cropped based on individual species-specific distributions (hereon SSA) from literature reviews (Table S1), IUCN polygons or ‘ranges’ (IUCN, 2020) and the terrestrial ecoregions where they occur. For creating the extent, we downloaded the current IUCN range maps for each species, then intersected those on ecoregions (Olson et al., 2001), then combined the results with selected ecoregions based on biogeographic knowledge of the species distributions and habitat preference from the literature reviews. For example, gaur habitat typically contains moist evergreen, semi-evergreen, and dry evergreen forests (Steinmetz et al., 2008; Tanasarnpaiboon, 2016), so we included these regions in our accessible areas. Further details on ecoregions included in accessible areas are in supplementary material, Table S4. To reduce overprediction and make our predictions closer to realised niche estimates, we used an occurrences-based threshold (OBR) method with ensemble models from (Mendes et al., 2020) for creating the spatially restricted ENM (hereon MSDM). OBR is an *a posteriori* method that restricts the suitable areas of our final ensemble models based on presence and the largest nearest neighbour distance among pairs of occurrences. Overall, we built four combinations between two accessible areas with and without MSDM methods for each species, including 1) No MSDM-SSA; 2) No MSDM-LA; 3) MSDM-SSA and 4) MSDM-LA.

### Model building

We processed the species occurrence files and environmental datasets in R 4.0.1 (R Core Team, 2020). We developed reproducible ecological niche models with optimized processing times using the ENMTML package (Andrade et al., 2020), following three main steps: 1) pre-processing, 2) processing and 3) post-processing.

In pre-processing, we performed occurrence thinning using 2 times the cell-size (1 km^2^) (Velazco et al., 2019) to reduce clustering of species records and sampling bias. We used principal component (PC) analysis (PCA) to reduce the collinearity of the predictors. We assigned species’ accessible areas to determine the species’ distributions using a mask function. We used random sampling to create pseudo-absence background points in a 1:1 ratio with presence points (Barbet-Massin et al., 2012). The occurrence and pseudo-absence data was divided into two sets for fitting the model (75%) and evaluating the fitted models (25%), using the bootstrapping partition method with 10 replications for each algorithm.

In the processing step, eight algorithms were used to build the ENMs, namely: BIOCLIM (Booth et al., 2014), Generalized Linear Models (McCullagh & Nelder, 1989), Generalized Additive Models (Hastie, 2018), Random Forest (Liaw & Wiener, 2002), Support Vector Machine (Karatzoglou et al., 2004), Maximum Entropy default (Phillips et al., 2006), Maximum Likelihood (Royle et al., 2012) and Bayesian Gaussian Process (Golding, 2014). All models used the default settings from the ENMTML package, which included the functions from different packages (e.g. dismo, maxnet) based on the algorithms that use to fit the models. The data type used for each algorithm is in supplementary materials, Table S5.

In the post-processing step, we created ensemble models using the weighted average (WMEAN) method based on the True Skill Statistic (TSS) values for building final habitat suitability and binary maps. The benefits of ensemble models are 1) robust decision-making (Ahmad et al., 2020); 2) reducing uncertainty (Marmion et al., 2009); and 3) a combination of several models into one model prediction (Kindt, 2018). We used TSS to calculate threshold values to convert habitat suitability maps into binary suitability maps (0 = unsuitable and 1 = suitable). We used TSS and area under the curve (AUC) for evaluating model performance. The TSS threshold is calculated using the maximum summed specificity and sensitivity and is not based on prevalence, where an equal TSS score for given models means similar performance (Allouche et al., 2006). Therefore, we selected the final models from the best TSS of weighted average ensemble models. We assessed the model’s accuracy by plotting a new dataset of species occurrences obtained from camera traps and human observations (https://www.gbif.org/) on the binary maps. Because, for example, gaur have been recorded to walk up to 6.3 km a day (mean 1.6 km (Rizal et al., 2020)), we created a 5 km buffer zone measured from the edges of the suitable pixels to include occurrences within the travel distance of wild bovids’ movement (Ahrestani & Karanth, 2014; Gardner et al., 2014). The percentage of points inside and outside the suitable areas and the buffer zone was calculated for each species. We present all the results, then only models with high prediction accuracy (greater than 80%,(Zhang et al., 2015)) are selected for further analyses. The total suitable areas of the best TSS binary map models were calculated using the zonal function in the raster R package (Hijmans, 2023). Then, we summed the pixels of the best TSS binary maps to generate the map of species number.

### Protected area analyses

The source for our protected areas map was the World Database of Protected Area (WDPA) (UNEP-WCMC & IUCN, 2021). We classified protected areas based on IUCN protected areas from WPDA into 8 categories, including categories 1 to 6 as IUCN management categories I to VI; category 7 as ‘not applicable’, which includes ‘not reported’, ‘not applicable’ and ‘not assigned’ protected areas; and category 8 as non-protected areas, which are the remaining areas that have not been classified as IUCN categories 1 to 7 (UNEP-WCMC and IUCN, 2021). Then, we used the zonal function in the Raster package to calculate overlapping areas between the suitable areas and protected areas for each species.

We calculated the percentage of suitable areas in WDPA polygons using the exact_extract function in exactextractr package (Baston et al., 2021) for extracting the suitable areas (values = 1) from binary map rasters in each WDPA polygon. Then, we classified each PA into 5 different suitability categories based on the percentage of suitable habitat in the PA: low suitability (0 - 20%); low - medium suitability (>20 - 40%); medium suitability (>40 - 60%); high suitability (>60 - 80%) and very high suitability (≥80%), and selected only the PAs that have the proportion of suitable area larger than species home range in the result. We have provided the code for creating the models in a GitHub repository.

## Results

We compiled 33,664 occurrence records. After filtering and spatial thinning, we used 3,708 points for modelling: 2,732 for gaur, 361 for banteng, 92 for wild water buffalo, 388 for mainland serow, and 135 for Chinese goral. The majority of the thinning occurrences (77%) were collected in Thailand, India and other countries in mainland SEA; see Table 1**Error! Reference source not found**. for details on the data coverage by country and supplementary materials Table S10 for details on the study sites.

We found that the PCA reduced the 28 environmental variables into 12 PCs that explained 95% of the environmental variance in the variables for the LA models for all species. The PCs for SSA models explained more than 96% of the total variance and the PC number varied by species, comprising 13 PCs (wild water buffalo), 11 PCs (gaur, mainland serow), and 10 PCs (banteng, Chinese goral). The bioclimatic variables were important variables in all species models. For LA models, the first two axes (PC1 and PC2) have high contributions from the annual mean temperature (bio01), mean temperature of the coldest month (bio06), mean temperature of the driest quarter (bio09) and mean temperature of the warmest quarter (bio10). The first two axes of SSA models showed high positive contributions from mean temperature of the coldest month (gaur), minimum temperature of coldest month (banteng, mainland serow), annual mean temperature (wild water buffalo, mainland serow), and precipitation of the wettest quarter (Chinese goral). We also found that NDVI, elevation, slope and human population density have less effect on explaining the variability for the first two PCs for all species. The correlations between PCs and individual environmental variables, PC biplots and percentage of explained variance are summarised in supplementary materials, Table S6 and Figure S2.

### Ecological niche models

Overall, all ensemble models showed high performance both for TSS and the area under the curve (AUC) with the highest performing models over 0.8 for all species (Table 2). Models with species-specific accessible areas were not always the best performing models, but most ensemble models performed above 0.7 TSS. The habitat suitability prediction maps using the best model ensembles are in supplementary materials, Figure S3 (SSA) Figure S4 (LA), Figure S5 (selected the best model of SSA and LA) and the binary maps which were used for calculating the suitable area in Figure S6. The performance of spatially restricted ensembles was higher in comparison with the No MSDM models, as the TSS was improved for banteng, Chinese goral and wild water buffalo. The lowest performing model for wild water buffalo was the No MSDM-SSA (TSS = 0.57). The best model for gaur was No MSDM-LA, banteng and Chinese goral is MSDM-LA, wild water buffalo is MSDM-SSA, and mainland serow is No MSDM-SSA. We found that all species have small predicted suitable habitats. Moreover, all species models predicted less than 50% of the suitable areas inside PAs.

**Table 2.**
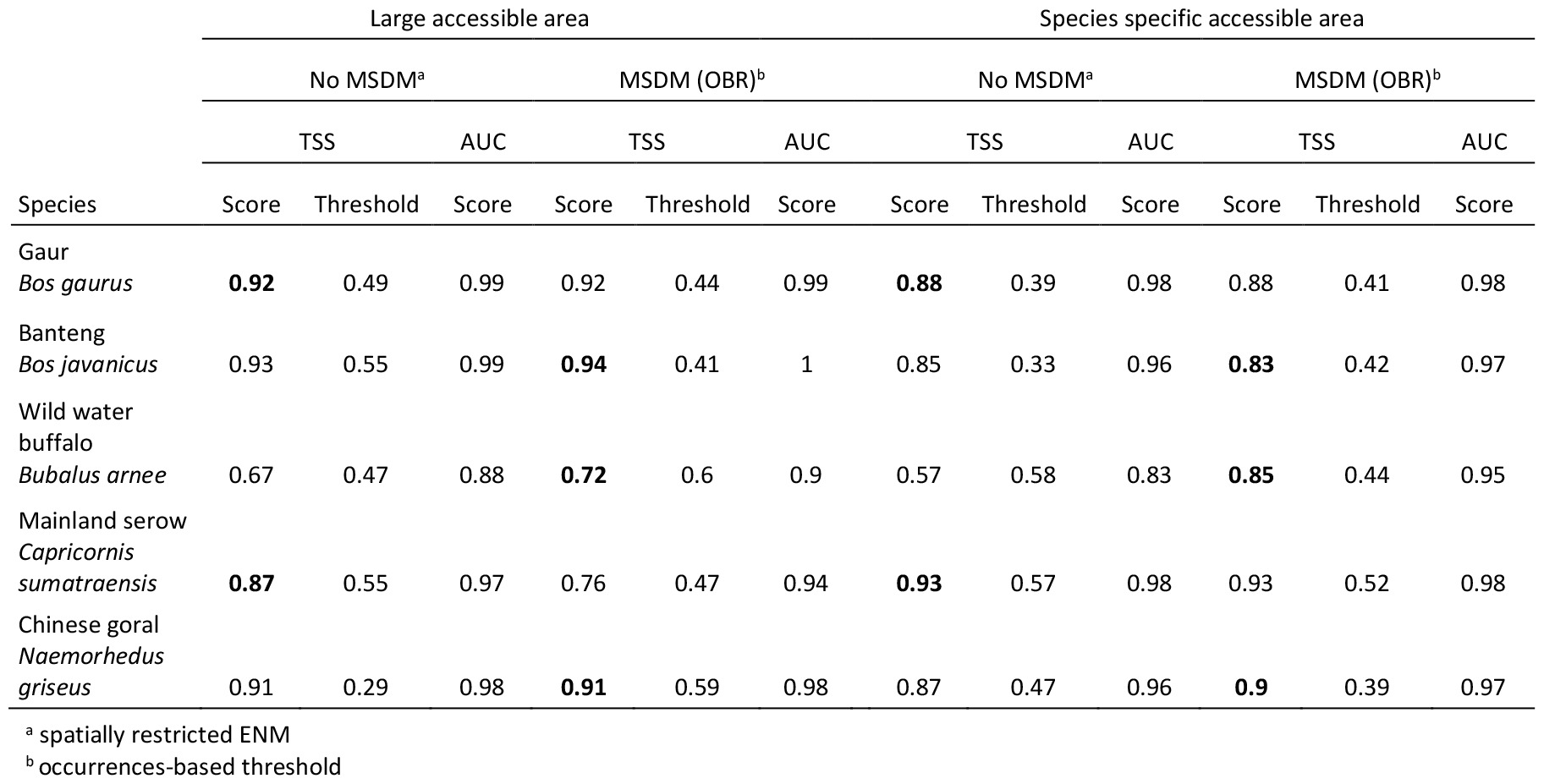
True Skill Statistics (TSS) and Area Under the Curve (AUC) values of the weighted average ensemble, and the threshold values for binary maps for five species classified by accessible area type and MSDM method.Best performing models for each accessible area by TSS are shown in **Boldface**.

The total of the suitable areas in km^2^ for each species and country are shown in Figure 2 and Suitable areas calculated from the best model are in supplementary materials Table S8 and the IUCN protected areas for all types of models are in Figure S7.

**Figure 2.**
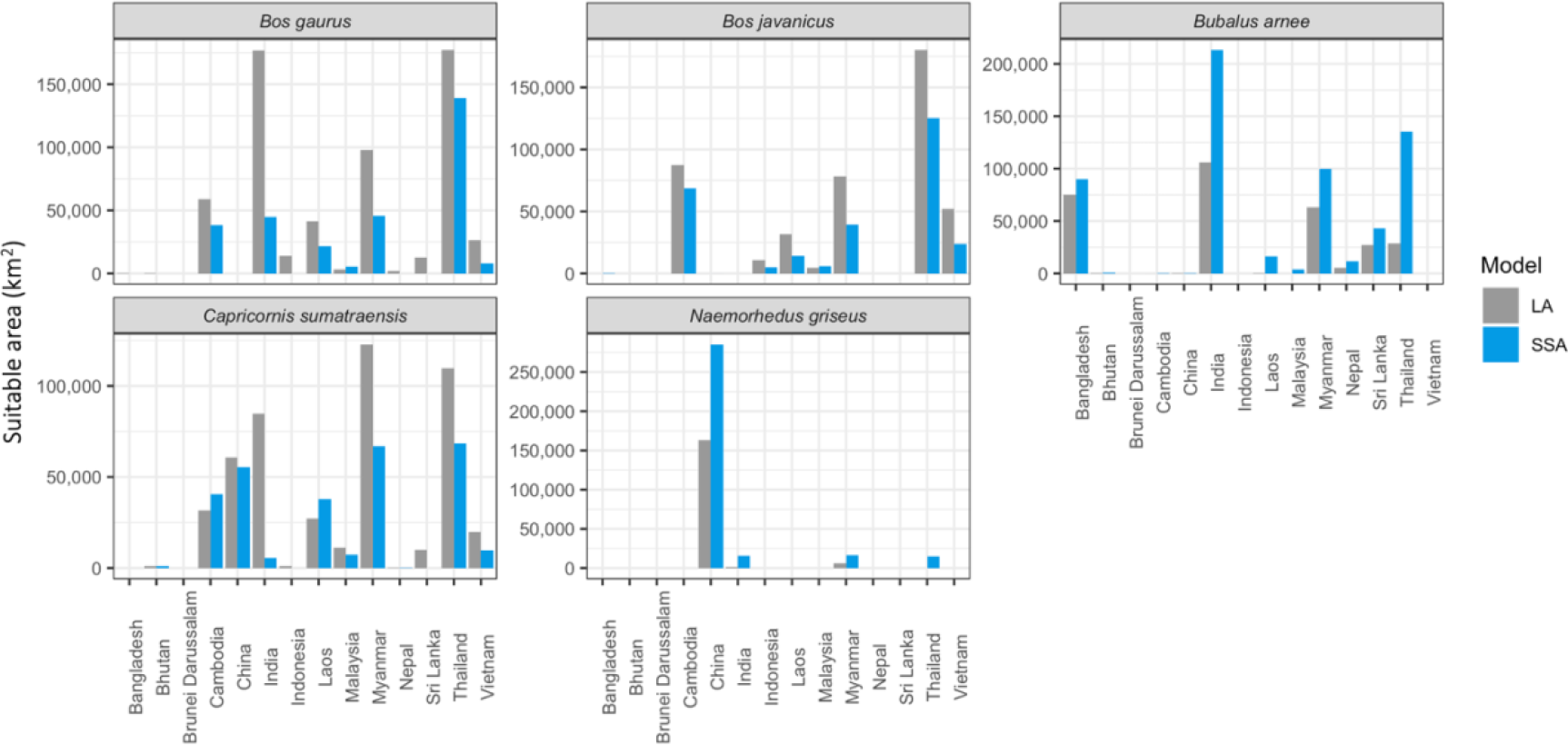
Total of the suitable area in km^2^ for each species and countries. Blue is the species-specific accessible area (SSA) and grey is the large accessible area models (LA) (see details in the supplementary Table S7).

Our model’s out-of-sample predictions with new species occurrences demonstrated a higher prediction accuracy within Thailand than the entire distribution, and this was further improved by including 5 km buffer zones, with the exception of Chinese goral, which exhibited poor accuracy across all scales (Table 3 and Figure 3). Implementing a buffer zone improves the accuracy of all four remaining species. For large herbivore species gaur, banteng and wild water buffalo, the model cropped to Thailand showed a higher accuracy (>80%) compared to the entire distribution (∼60-80%). We selected only model predictions with a high accuracy percentage, greater than 80%, for further analyses. As a result, three species, including gaur, banteng, and wild water buffalo, were retained, while two species, mainland serow and Chinese goral, were excluded from the rest of the study. Furthermore, we cropped the entire distribution to focus only on the result within Thailand as the number of data collection and model predictions is higher compared to the entire species distribution. The result of the entire distribution for all species can be found in the supplementary material, Figure S3 and Figure S4.

**Table 3.**
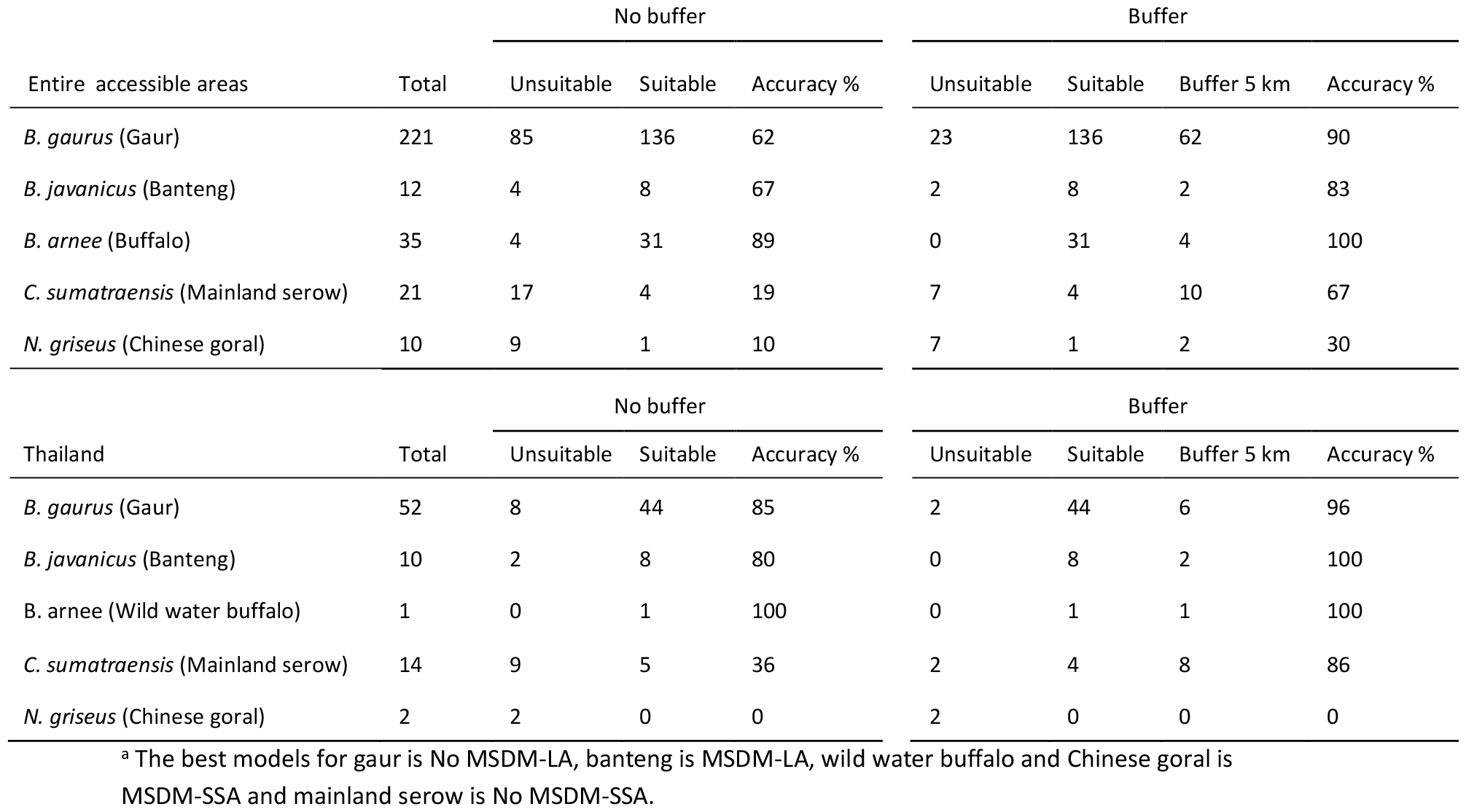
Comparison of the accuracy of the selected best models^a^ in predicting out-of-sample data for the entire accessible areas range and Thailand.

| Entire accessible areas | Total | No buffer |  |  | Buffer |  |  |  |
| --- | --- | --- | --- | --- | --- | --- | --- | --- |
|  |  | Unsuitable | Suitable | Accuracy % | Unsuitable | Suitable | Buffer 5 km | Accuracy % |
| *B. gaurus* (Gaur) | 221 | 85 | 136 | 62 | 23 | 136 | 62 | 90 |
| *B. javanicus* (Banteng) | 12 | 4 | 8 | 67 | 2 | 8 | 2 | 83 |
| *B. arnee* (Buffalo) | 35 | 4 | 31 | 89 | 0 | 31 | 4 | 100 |
| *C. sumatraensis* (Mainland serow) | 21 | 17 | 4 | 19 | 7 | 4 | 10 | 67 |
| *N. griseus* (Chinese goral) | 10 | 9 | 1 | 10 | 7 | 1 | 2 | 30 |

| Thailand | Total | No buffer |  |  | Buffer |  |  |  |
| --- | --- | --- | --- | --- | --- | --- | --- | --- |
|  |  | Unsuitable | Suitable | Accuracy % | Unsuitable | Suitable | Buffer 5 km | Accuracy % |
| *B. gaurus* (Gaur) | 52 | 8 | 44 | 85 | 2 | 44 | 6 | 96 |
| *B. javanicus* (Banteng) | 10 | 2 | 8 | 80 | 0 | 8 | 2 | 100 |
| *B. arnee* (Wild water buffalo) | 1 | 0 | 1 | 100 | 0 | 1 | 1 | 100 |
| *C. sumatraensis* (Mainland serow) | 14 | 9 | 5 | 36 | 2 | 4 | 8 | 86 |
| *N. griseus* (Chinese goral) | 2 | 2 | 0 | 0 | 2 | 0 | 0 | 0 |

<sup>a</sup> The best models for gaur is No MSDM-LA, banteng is MSDM-LA, wild water buffalo and Chinese goral is MSDM-SSA and mainland serow is No MSDM-SSA.

**Figure 3.**
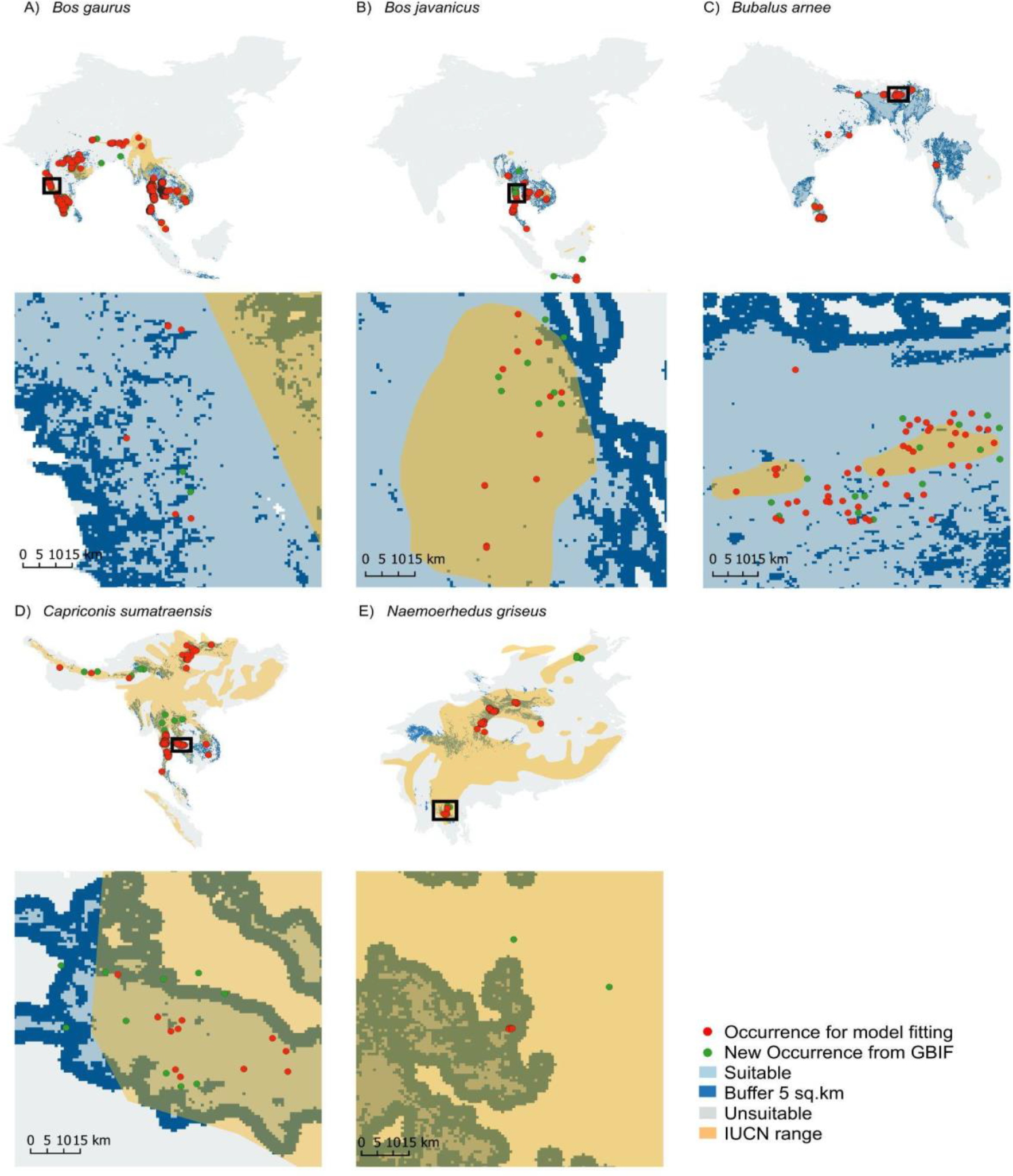
Model prediction testing for five bovid species (A-E) by calculating the percentage of the out of sample points that fall inside the model predicted suitable areas (blue). The model fitting datasets (red) were mainly within the suitable areas compared to the new occurrence dataset (green). IUCN ranges show greater areas than the predictions for mainland serow and Chinese goral. Some of the occurrence data were distributed outside both the model predicted suitable area and IUCN range.

Nearest distance from out of sample points to suitable area

| Species | point | Distance (km) |  |  |
| --- | --- | --- | --- | --- |
|  |  | Min | Mean | Max |
| Gaur | 52 | 0.0047 | 1.54 | 22.4 |
| Banteng | 10 | 0.0323 | 4.72 | 39.9 |
| Wild water buffalo | 1 | 0.811 | 0.811 | 0.811 |
| Mainland Serow | 14 | 0.00668 | 6.07 | 38.1 |
| Chinese goral | 2 | 0.147 | 1.54 | 2.93 |

### Identifying priority areas for conservation

Most suitable habitats in protected areas are located in IUCN category Ia (Strict nature reserve), Ib (Wilderness area) and II (National Park) areas for the best TSS models for all species, while IUCN category V (Protected landscape or seascape) has the least. Overall, more than half of the species’ suitable habitat is not under any form of protection defined by the WDPA (supplementary materials, Table S8, Figure S8). The proportion of the suitable area in each WDPA of the best models from SSA and LA for each species are presented in supplementary materials, Figure S9 and Figure 10.

In Thailand, we identified a high percentage (≥80%) of suitable area of Thailand for gaur in 122 PAs (74,268 km^2^; 15% of Thailand), banteng in 102 PAs (59,528 km^2^; 12% of Thailand), and wild water buffalo in 3 PAs (559 km^2^; 0.1 % of Thailand). A high proportion of the suitable area for gaur and banteng is in Thungyai Naresuan, Kaengkrachan and Huai Kha Khaeng, and for wild water buffalo in Phu Wua WS and Dong Yai WS in eastern DPKY-FC (Figure 4 and Figure 5). The hotspots for five species can be found in supplementary materials, Figure S11.

**Figure 4.**
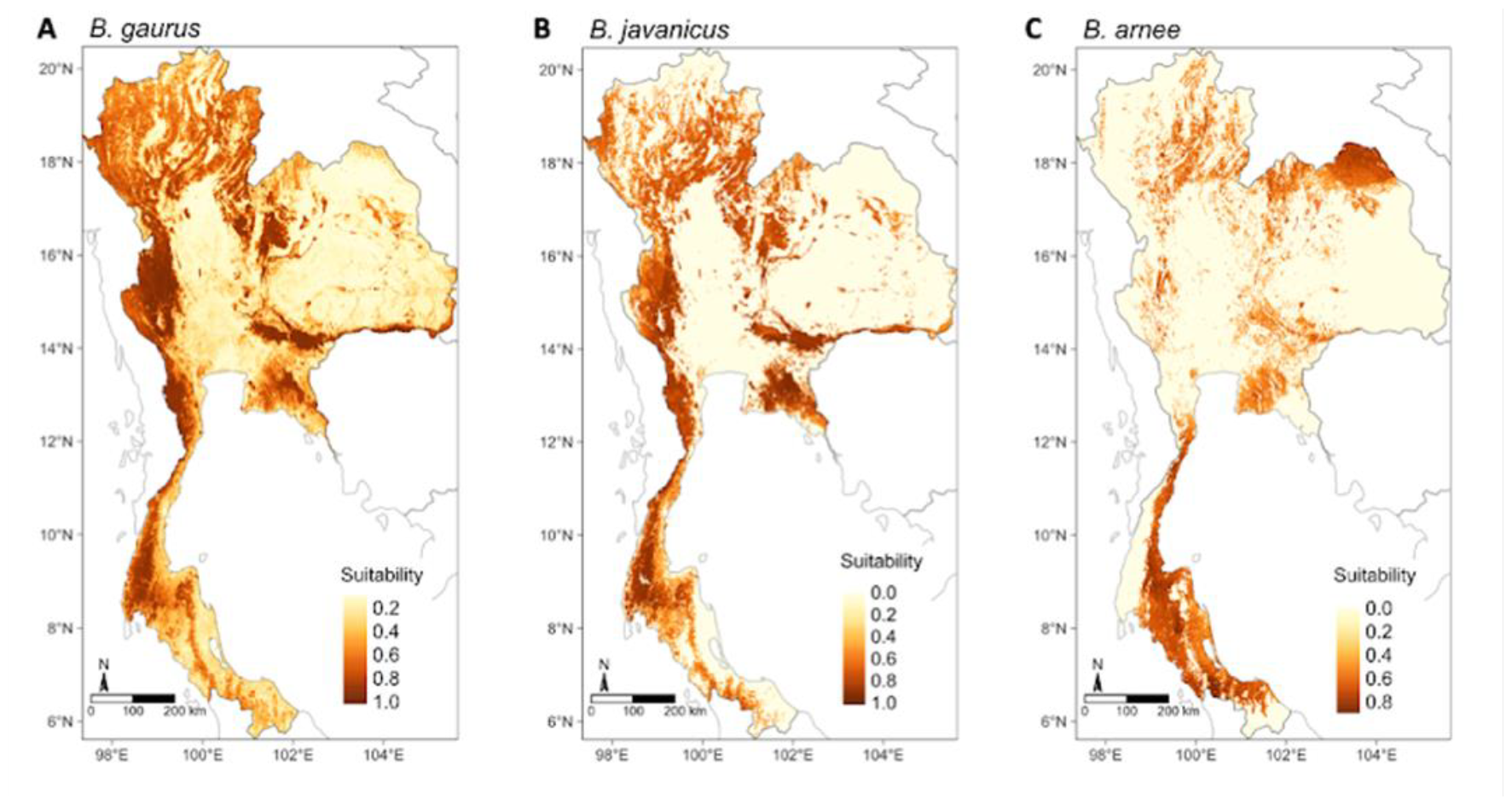
Habitat suitability prediction maps of three wild bovids species in Thailand: gaur (*B. gaurus*), banteng (*B. javanicus*) and wild water buffalo (*B. arnee*) species (A-C) using the best model from the weighted average ensemble. The value ranges from 0-1: yellow represents low suitability and dark brown represents high suitability. Interactive maps are provided in the online supplementary material (link).

**Figure 5.**
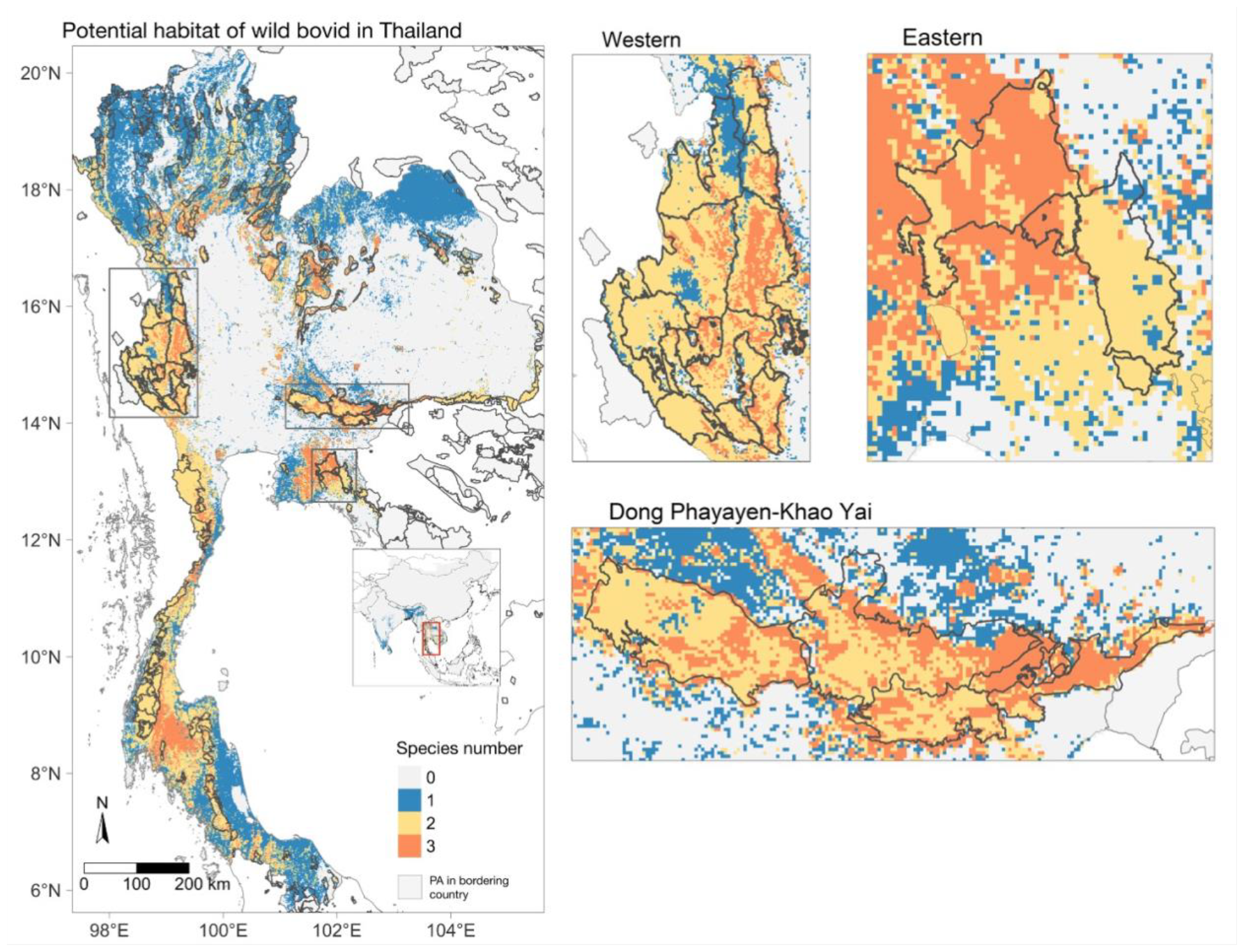
Estimated species richness of three wild bovids in Thailand. The species are gaur, banteng, and wild water buffalo. Frames A-C focus on (A) Western Forest Complex (WEFCOM), (B) Dong Phayayen-Khao Yai Forest Complex (DPKY-FC) and (C) Eastern Forest Complex, where the overlapping suitable areas of all species (n=3). Western, Dong Phayayen-Khao Yai and Eastern forests have suitable areas for gaur, banteng and wild water buffalo for both inside PAs and also in the surrounding areas.

Proportions range from 0 (all unsuitable) to 1 (all suitable), with suitability determined by thresholds from species best performing models. (A) gaur (*Bos gaurus*), (B) banteng (*Bos javanicus*), (C) wild water buffalo (*Bubalus arnee*).

We found that the highest percentage of suitable area was comprised of mixed deciduous forest for all species, followed by evergreen forest for gaur and banteng, and dry dipterocarp forest for wild water buffalo. We found a percentage of non-forest areas identified from the total suitable for all species: wild water buffalo (71%), banteng (33%), and gaur (24%). For more details of forest types by suitable areas, see Table 4 and Figure S12.

**Table 4.**
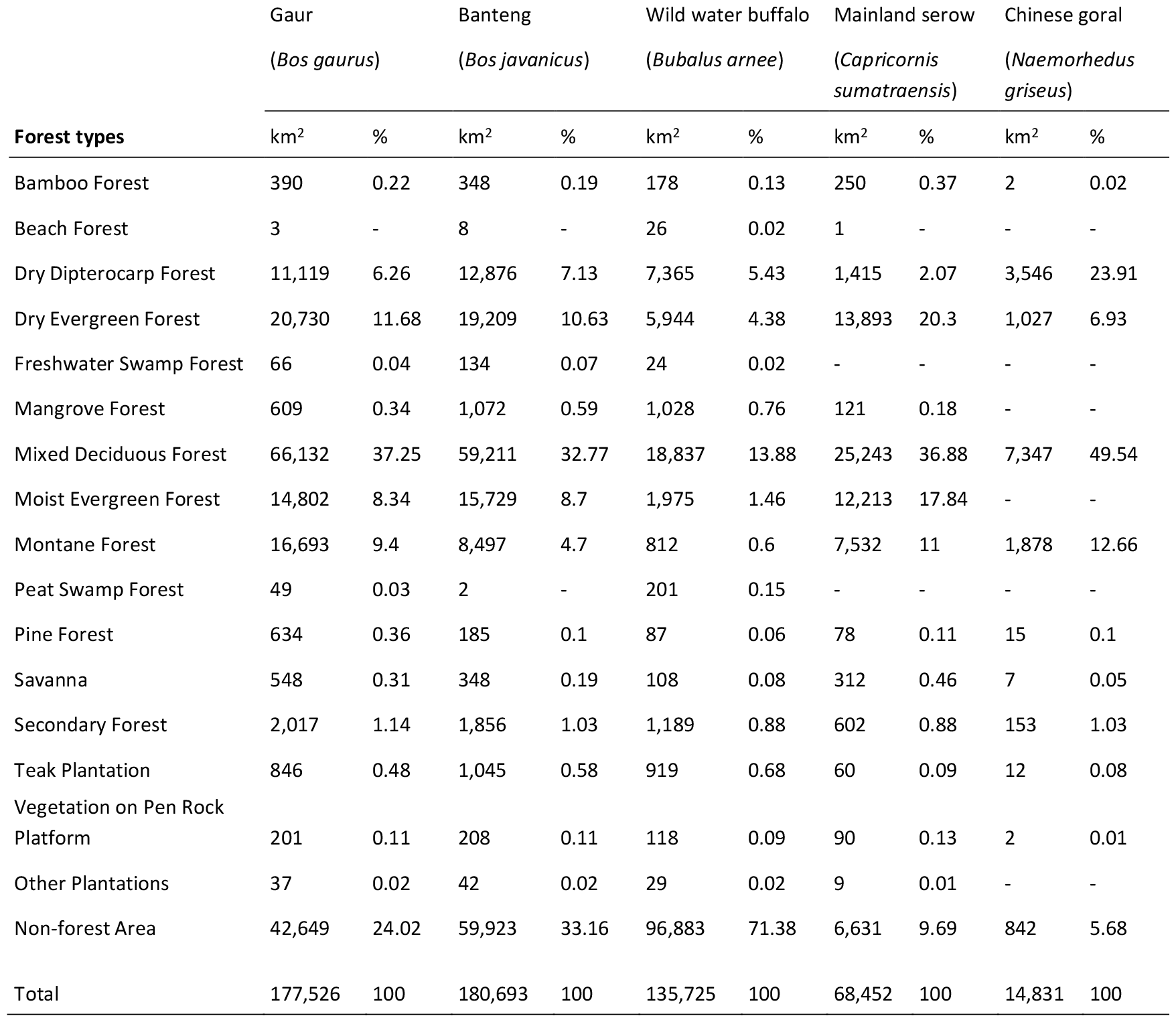
The suitable areas of five bovid species classified by forest types in Thailand.

## Discussion

We modelled the potential distribution for the five threatened wild bovid species, distributed in East, South and Southeast Asia. Our aim was to build predictive models to identify conservation areas, and potential species richness maps in their entire geographical distribution. However, the model predictions were better for Thailand, where most data were from for all but Chinese goal (Table 3), therefore we were focus on Thailand. We found our models were able to predict the presence of out of sample observations well for three species, gaur, banteng, and wild water buffalo throughout the entire distribution (≥62%), but not mainland serow or Chinese goral (≤19%). We identified that suitable areas were fragmented and often (50%) located outside PAs. Those suitable areas outside PAs could possibly be managed as corridors or buffer zones to connect currently fragmented bovid populations, thereby enhancing long-term wild bovid conservation success (Karanth, 2016; Penjor et al., 2021) which requires further investigations. For example, a corridor was built within DPKY-FC and showed the possibilities of connecting the western forest complex and Kaengkrachan NP to conserve the endangered tiger population (Sukmasuang et al., 2020; Suttidate et al., 2021).

Our study found that most suitable areas for gaur were similar to IUCN range assessments (Duckworth et al., 2016) and consistent with studies that have confirmed species presences, such as in Thailand’s PAs (Prayoon et al., 2021), Myanmar (Hein et al., 2020) and Western Ghats in southwestern India and Manas WS in the Himalayan foothills (Choudhury, 2002). However, there are differences. Our study predicted larger gaur suitable habitats in Thailand inside (∼82,400 km^2^) and outside (95,000 km^2^) PAs than Prayoon et al. (2021), who predicted 39,508 km^2^ of total suitable habitat. Choudhury (2002) predicted their distributions in Western Ghats, Central and North-eastern India which included larger than our predictions. Our predictions used NDVI and land coverage fractions (Table S3) for predicting greenness, which may be useful for predicting the vegetation quality and availability for ungulates (Borowik et al., 2013). However, NDVI is difficult to differentiate vegetation variations (Didan et al., 2015; Martinez & Labib, 2023), such as between specific agricultural areas, grassland, and dense forest canopy. This may include other vegetation types other than the species’ habitat in suitable areas and estimated larger suitable areas predicted in non-forest areas and non-PAs to be identified in our study, compared to Prayoon et al’s study. Other studies suggest that gaur does use crop plantations or man-made grasslands, which may increase the suitable areas in our prediction, even if these are not their natural habitats and lead to conflict between humans and gaur (Chaiyarat et al., 2021).

Our best model predicted larger suitable areas (446,075 km^2^) for banteng than the IUCN-SSC report released in 2010 (∼ 209,000 km^2^) (IUCN-SSC AWCS Group, 2010). We found a high percentage of predicted suitable areas in Eastern Plains Landscape (ELP) and Chhaeb WS in Cambodia; the former supports the likely largest banteng population globally (Gray et al., 2012). However, our results showed low habitat suitability in Sundaic Southeast Asia, with just 2% of the total suitable area in Indonesia (mainly in Alas Purwo NP, Java) and 2% of the total suitable area in Malaysia. Banteng populations and habitats in Southeast Asian islands (Borneo, Java, and Bali) are threatened due to hunting for horn and meat consumption and habitat loss (Dewi et al., 2020). In Thailand, we found high suitability similar to previous studies in Eastern (Menkham et al., 2019) and Western forest complexes (Jornburom et al., 2020), including reintroduction areas in Salak Pra WS (Chaiyarat et al., 2019) and where recent recolonisation by natural population movement has occurred in Mae Wong NP (Phoonjampa et al., 2021).

Wild water buffalo has been domesticated and bred as livestock, making it hard to distinguish between the free-grazing domestic buffalo and wild water buffalo as domesticated animals may replace wild animals in suitable habitats and cause the high suitable area prediction outside PAs, especially in overlapping habitats (Zhang et al., 2020). We estimate the highest percentages of suitable areas at Kaziranga NP in India, currently with the largest population of wild water buffalo (Kaul et al., 2019). Grasslands and flood plain areas of Manas NP (500 km^2^) and Kaziranga NP (>850 km^2^) in India contain the most suitable habitat and are the main population strongholds for wild water buffalo (Choudhury, 2014). In Thailand, this type of habitat can be found in many places, but it is not often represented in protected areas. Wild water buffalo are only found in Huai Kha Kheang WS parts of the Western Forest Complex. Our model predicts that only 43% of Huai Kha Kheang Wildlife Sanctuary is suitable for this species, primarily because the floodplains are mainly situated close to the mainstream in the middle of the PA. Additionally, the population has remained constant for decades, which could be attributed to a single population group or constraints within suitable habitats.

The three selected species showed overlapping suitable areas in the Western Forest Complex, Eastern Forest Complex, and Dong Phayayen-Khao Yai Forest Complexes (DPKY-FC). These forest complexes encompass extensive areas of high wildlife biodiversity and diverse forest types, including several contiguous protected areas (PAs) situated at the borders of Cambodia and Myanmar. The Western Forest Complex is the largest conservation area in Thailand where these wild bovids still exist, while the DPKY-FC maintains a higher population of gaur as they are mainly covered by evergreen forest. The Eastern Forest Complex sustains a large population of banteng because most of the main vegetation consists of deciduous and dipterocarp forest. Gaur uses a diversity of types of habitats and prefers denser canopy at higher elevation than banteng, which tends to inhabit in dry and open habitats such as dry dipterocarp and deciduous forests (Gray & Phan, 2011; Steinmetz, 2004). Wild water buffalo also shares overlapping areas with these two species, despite its distribution being found exclusively in Huai Kha Khaeng Wildlife Sanctuary. We recommend protecting these important suitable habitats to ensure the protection of wild bovids. This may involve implementing active patrolling to reduce illegal intrusions, snare removal and habitat management based on their diet diversity (McShea et al., 2019). Additionally, one option to maintain wild water buffalo populations is to reintroduce them into their historical range, from which they have been extirpated. This method could be evaluated by combining predicted suitable areas with several important factors such as vegetation types, forage biomass, carrying capacity and hunting pressure (Bora et al., 2024).

In this study, we included all subspecies data points in our model ensembles as we aim to extrapolate and predict the entire range of species’ habitat suitability, but this may increase uncertainty (Dormann, 2007). These five bovids have multiple subspecies, including 3 subspecies of gaur (Duckworth et al., 2016), banteng (Gardner et al., 2016) wild water buffalo (Kaul et al., 2019) and mainland serow (Mori et al., 2019), and 2 subspecies of Chinese goral (Duckworth et al., 2008). Subspecies may vary in niche, climate and biological interactions that could affect the model predictions. The low habitat suitability of our study in Borneo for banteng could be because climatic and geographic conditions differ for *B*.*j. lowi* compared to those in mainland Asia, affecting model transferability across different regions (Zhu et al., 2021). Equally, Mori et al. (2019) suggest that Chinese goral (*N. griseus*) should be reclassified within Brown goral (*N. goral*) together and Burmese goral (*N. evansi*) that together with *N. griseus* should be split to become an individual species. Future analyses must consider these taxonomic reclassifications. However, we modelled species level habitat suitability, rather than the subspecies, as we assume that there is less likely to be habitat and environmental condition variation at the subspecies level for these bovids (Smith et al., 2019).

We found that using the MSDM OBR technique showed a better predicted suitable area of the ecological niche, closer to the real distribution for species with more restricted ranges like banteng, wild water buffalo and Chinese goral, with higher performance TSS values compared to No MSDM models. We recommend restricting the accessible area for predicting wild water buffalo potential habitat to reduce overprediction caused by overlapping areas with domestic water buffalo.

We also used ensemble approaches, to obtain better predictive performance than from any single model type, but further analyses could also look at individual model results using different parameters, such as differing pseudo-absence background point ratios. The equal ratio of presence to pseudo-absence (1:1 ratio) has been used in several types of model like general linear models, artificial neural networks, and Maxent models, and it is also recommended for use in ensemble models when dealing with small sample sizes (Liu et al., 2019).

### Limitations

We acknowledge sampling deficiencies across the regions. We had fewer occurrences in Vietnam, Laos, Myanmar and Indonesia compared to Thailand, from which a large number of our data points came (30,512 points in Thailand, 3,152 points outside Thailand, **Error! Reference source not found**.). Occurrence data based on data accessibility may have sampling bias, particularly with clustered points for gaur, banteng, and mainland serow. We minimised these biases through spatial thinning (Aiello-Lammens et al., 2015). Since we found large amounts of suitable areas outside of Thailand, we suggest that future studies should focus on monitoring bovid populations in other countries, especially in India and Myanmar. However, because of this and the model performance, we focused on Thailand.

Missing data data has impacted some results. The model TSS values for endangered banteng and Chinese goral are over 0.8, yet our models predict unsuitable areas in part of Indonesia (east and central Kalimantan; Dewi et al. 2020) for banteng and China (e.g. Beijing and northeast Inner Mongolia; (Yang et al., 2019) for Chinese goral from which these species have been reported. This would likely be improved if more spatial data were available for these species.

We used a new dataset of species occurrences to assess our model’s performance with a 5 km buffer zone, aiming to enhance modelling accuracy. Given these species have quite large home ranges and daily movements, adding a buffer to represent this movement unsurprisingly lead to better model predictions for all species, but most notably for mainland serow, changing the out of sample prediction from 19% to 67% for the entire region and 36% to 86% for Thailand. The buffer zone may indicate the utilisation of unsuitable areas of the species near forested regions, such as secondary forests, agricultural areas, or water resources, which possibly extend these buffer areas from the protected aeras to enhance the wildlife protection.

The spatial restriction method, OBR, can be sensitive to the distribution of occurrence data, because it keeps predicted suitable areas close to the occurrence locations. This may lead to the exclusion of potential suitable areas driven by a lack of occurrence data in those areas. For example, the wild water buffalo No MSDM predicted potentially suitable habitat around the Sre Pok Wildlife Sanctuary in Cambodia where the species is distributed (Gray et al., 2012), but after the MSDM, this potential habitat was excluded as we lack occurrence data in Cambodia. Although our study showed slightly different TSS values between two different accessible area extents, we encourage testing the different accessible areas as it affects the model results (Anderson & Raza, 2010). Moreover, model performance varied with accessible area sizes and spatial restrictions, emphasising the need for careful accessible area definition in ecological modelling (Barve et al., 2011). Further, future analyses may try to better account for the current presence of species by accounting for factors such as hunting using other proxies, such as other human-disturbance metrics like distance from roads (Lim et al., 2021).

## Conclusion

Our study provided an overview of the suitable remaining habitat for threatened bovid species at a regional scale using high-resolution environmental variables and species occurrence data from multiple observation methods. Our predictions showed that the suitable areas are small and fragmented for all species, and more than 50% of suitable areas are outside of protected areas. Those suitable areas outside PAs could possibly become efficient conservation areas, such as forest corridors or buffer zones to connect fragmented bovid populations and enhance long-term habitat conservation. Our predictions may inform conservation actions to avoid further defaunation of wild bovidae such as the management of human-wildlife conflicts and habitat quality for long-term species survival.

## Acknowledgements

We thank all the data contributions and collaborations from these institutions:

Dr Supagit Vinitpornsawan (Director of Wildlife Conservation Area Management and Education Center, Wildlife Conservation Office, Department of National Parks, Wildlife and Plant Conservation, Thailand) for animal track and sign data in Thailand. For camera trap data in Thailand: Naret Seuaturien (WWF Thailand), Manoon Pliosungnoen, Department of Environmental and Forest Biology, State University of New York College of Environmental Science), Wanlop Chutipong (Conservation Ecology Program, Pilot Plant Development and Training Institute, King Mongkut’s University of Technology Thonburi), Nucharin Songsasen (Smithsonian’s National Zoo and Conservation Biology Institute), Lon I. Grassman, Jr. (Feline Research Program, Caesar Kleberg Wildlife Research Institute, Texas A&M University, USA), Freeland Foundation, WCS Thailand, Smithsonian Institution, PANTHERA USA in Thailand.

WWF Thailand would like to thank: WWF Germany, WWF Sweden, WWF US, B.Grimm, WWF Japan, and WWF Switzerland for wonderful support of field projects, and Department of National Parks, Wildlife, and Plant Conservation for kind permission and collaboration. WCS Cambodia, Friends of Wildlife, Wildlife Alliance and Ministry of Environment, Royal Government of Cambodia for the camera trap data in Cambodia. CarBi Project of WWF Lao and WWF Greater Mekong for the camera trap data in Laos. Camera trap in Myanmar: Friends of Wildlife and Data collecting in Tanintharyi, Myanmar was partially funded by the European Union, Helmsley Charitable Trust and mainly funded by Integrated Tiger Habitat Conservation Project, through the Fauna & Flora International (FFI). Open source databases: https://www.gbif.org, camera trap data from eMammals (https://emammal.si.edu/) by William J. McShea (Conservation Ecology Center, Smithsonian Conservation Biology Institute) and Megan Baker-Whatton (Smithsonian Conservation Biology Institute, and George Mason University) under the project: Qionglai Mountains Project, Liangshan Mountains Project, Habitat Connectivity in Minshan Mountains, Qinling Project, HKK ForestGEO Project, and Carnivore Intraguild Interactions in Select Thailand Reserves. Also, the other researchers who share the species occurrence data with us. We thank the IUCN specialists for commenting on the early version of the habitat suitability maps. The authors wish to acknowledge the use of New Zealand eScience Infrastructure (NeSI) high-performance computing facilities, consulting support and/or training services as part of this research.

## Funding Statement

WH was supported by Manaaki New Zealand Scholarships. DTSH, RLM, RSJ were supported by Bryce Carmine and Anne Carmine (née Percival), through the Massey University Foundation. DTSH was supported by the Percival Carmine Chair in Epidemiology and Public Health and Royal Society Te Apūrangi (grant no. RDF-MAU1701). EA supported by U.S. Fish & Wildlife Service Rhinoceros and Tiger Conservation Fund, David Shepherd Wildlife Foundation, Care for the Wild International/Born Free Foundation, Point Defiance Zoo & Aquarium and 21st Century Tiger. DN was supported by King Mongkut’s University of Technology Thonburi (grant no. WOR1-2557–2558); International Association for Bear Research and Management (Research & Conservation grant), 2012 & 2014 and The Asahi Glass Foundation (Research Grant), 2003. Antony Lynam was supported by TRF/BIOTEC Special Program for Biodiversity Research and Training. George A. Gale was supported by TRF/BIOTEC Special Program for Biodiversity Research and Training (BRT R 346001) and National Science and Technology Development Agency (grant no. NSTDA P-11-00390). Data collecting in Tanintharyi, Myanmar was partially funded by the European Union, Helmsley Charitable Trust and mainly funded by Integrated Tiger Habitat Conservation Project - grant no. ITHCP1338, through the Fauna & Flora International.

## Data Accessibility

Data available upon request

## Competing Interests

We have no competing interests.

## Authors’ Contributions

**Alex Riggio**: Investigation, Resources, Writing - Review and Editing.

**Alexander Godfrey**: Investigation, Resources, Writing - Review and Editing.

**Anony Lynam**: Investigation, Resources, Writing - Review and Editing

**David T. S. Hayman:** Conceptualization, Methodology, Validation, Supervision, Writing - Original Draft, Writing - review and Editing, Project administration, Funding acquisition.

**Dusit Ngoprasert**: Investigation, Resources, Writing - Review and Editing.

**Eric Ash**: Investigation, Resources, Writing - Review and Editing.

**Francesco Bisi**: Investigation, Resources, Writing - Review and Editing.

**George A. Gale**: Investigation, Resources.

**Giacomo Cremonesi**: Investigation, Resources, Writing - Review and Editing.

**Gopalasamy Reuben Clements**: Investigation, Resources, Writing - Review and Editing.

**Jonathan C Marshall**: Methodology, Software, Validation, Formal analysis Supervision, Writing - Original Draft, Review and Editing.

**Marnoch Yindee**: Investigation, Resources, Writing - Review and Editing.

**Nay Myo Shwe:** Investigation, Resources.

**Pin Chanratana**: Investigation, Resources.

**Reju Sam John**: Methodology, Software, Validation, Formal analysis, Supervision, Writing - Original Draft, Review and Editing.

**Renata L. Muylaert:** Methodology, Software, Validation, Formal analysis, Supervision, Writing - Original Draft, Review and Editing.

**Robert Steinmetz**, Investigation, Resources, Writing - Review and Editing.

**Rungnapa Phoonjampa**: Investigation, Resources.

**Saw Soe Aung**: Investigation, Resources.

**Seri Nakbun**: Investigation, Resources.

**Stephanie Schuttler**: Investigation, Resources, Writing - Review and Editing.

**Thomas N. E. Gray**: Investigation, Resources, Writing - Review and Editing.

**Wantida Horpiencharoen**: Conceptualization, Methodology, Software, Formal analysis, Data Curation, Writing - Original Draft, Writing - review and Editing, Project administration, Funding acquisition.

**Worrapan Phumanee**: Investigation, Resources.

All authors read the text, provided comments, suggestions and corrections, and approved the final version.

## Notes

https://github.com/Wantidah/ENM-Bovidae

### Competing Interest Statement

The authors have declared no competing interest.

### Summary of Updates

The method on selecting the models and maps was updated. Excluded the poor-performance models from the analysis.

