## Supplementary Materials for "Mapping threatened Thai bovids provides opportunities for improved conservation outcomes in Asia"

Contents


- Table S1 to S10
- Table S12 (link)
- Figure S1 to S11
- References

Table S1 Literature review summary prior to the selection of the accessible areas.

| Species | Study area | Study methods | Habitat suitability analysis method | References |
|---|---|---|---|---|
| Gaur *Bos gaurus* | India | Transect survey | Ecological niche factor analysis | Paliwal and Mathur (2012) |
| | | Direct observation, Animal signs surveys, | Distribution, population density | Choudhury (2002) |
| | | | Logistic regression | Imam and Kushwaha (2013) |
| | | Radio collar | Minimum Convex Polygon | Sankar et al. (2013) |
| | | Grid-based survey | Linear regression | Ahrestani et al. (2012) |
| | Nepal, Bhutan, Myanmar | Direct observation, Animal signs surveys | Distribution, population density | Choudhury (2002) |
| | Thailand | Secondary data, Field survey | Relative abundance | Srikosamatara and Suteethorn (1995), DNP (2010) |
| | | Animal signs survey | Distribution | Steinmetz (2004) |
| | | Animal signs survey | MAXENT | Trisurat et al. (2015), Planisong et al. (2019) |
| | | Direct observation, Camera trapping | Capture-recapture, Relative abundance | Tanasarnpaiboon (2016) |
| | | Transect survey | Animal density, Minimum convex polygon | Laichanthuek et al. (2017) |
| | Malaysia | Radio collar | Home range, Habitat use | Conry (1989) |
| | Laos | Animal signs survey | Descriptive | Steinmetz (2004) |
| | Vietnam | Secondary data, Direct observation, | Population density | Nguyen (2009) |
| | Myanmar | Animal signs survey | Distribution | Rabinowitz et al. (1995) |
| | Bhutan, Cambodia, China (Southern), India, Lao PDR, Malaysia (Peninsular), Myanmar, Nepal, Thailand, Vietnam | Review | | Ahrestani and Karanth (2014) |
| Gaur *Bos gaurus* | Bhutan, Cambodia, China (Southern), India, Lao PDR, Malaysia (Peninsular), Myanmar, Nepal, Thailand, Vietnam | IUCN assessment | - | Duckworth et al. (2016) |
| Banteng *Bos javanicus* | Thailand | Secondary data, Animal signs survey | Relative abundance | Srikosamatara and Suteethorn (1995) |
| | | Animal sign survey | Descriptive | Steinmetz (2004) |
| | | Field survey | Relative abundance, species richness | DNP (2010) |
| | | Animal signs survey | MAXENT | Trisurat et al. (2015); |
| | | Direct observation, Animal signs survey | Logistic regression model | Chaiyarat et al. (2017) |
| | | Radio collar | MAXENT | Chaiyarat et al. (2019) |
| | Vietnam | Secondary data, Direct observation | Population density | Nguyen (2009) |
| | | Animal signs survey | Occupancy | Pedrono et al. (2009) |

| Species | Study area | Study methods | Habitat suitability analysis method | References |
|---|---|---|---|---|
| | Cambodia | Camera trapping | Relative abundance | Gray and Phan (2011) |
| | Indonesia | Camera trapping | MAXENT | Rahman (2020) |
| | | Forest patrol | Multiple logistic regression | Imron et al. (2016) |
| | Malaysia | Camera trapping | MAXENT | Lim et al. (2021) |
| | Cambodia, Myanmar, Vietnam Malaysia (Sabah), Indonesia (Kalimantan, Jawa) | | | Gardner et al. (2014) |
| | Cambodia, Indonesia (Kalimantan, Jawa, Bali), Malaysia (Sabah), Myanmar, Thailand, Viet Nam | IUCN assessment | - | Gardner et al. (2016) |
| Wild water buffalo *Bubalus arnee* | Thailand | Transect survey | Logistic regression | Chaiyarat et al. (2004) |
| | | Field survey | Relative abundance, species richness | DNP (2010) |
| | | Transect survey | MAXENT | Khiowsree et al. (2015) |
| | Nepal | Population census | Population structure | Heinen and Kandel (2006) |
| | | Field survey | Intersection of habitat suitability criteria | Thapa et al. (2020) |
| | Sri Lanka | Population census | Population structure | Silva et al. (2012) |
| | Bhutan, Cambodia, India, Nepal, Thailand, Malaysia (Peninsular Malaysia), Sri Lanka | | | Choudhury (2014) |
| | Bhutan, Cambodia, India, Myanmar, Nepal, Thailand, Indonesia, Lao PDR, Malaysia (Peninsular Malaysia), Sri Lanka, Viet Nam | IUCN assessment | - | (Kaul et al., 2019) |
| Chinese goral *Naemorhedus griseus* | India | Transect survey, Interviews | Logistic regression | Singh and Kushwaha (2011) |
| | Thailand | Animal signs surveys, Direct observation | Home range, Population structure | Chaiyarat et al. (1999) |
| | | Field survey | Relative abundance, species richness | DNP (2010) |
| | | Radio collar | plotting occurrences | Buranapim et al. (2014) |
| | | Animal signs survey, | MAXENT, Multiple logistic regression model | Trisurat et al. (2015) |
| | China | Animal signs survey | Habitat use | Chen et al. (2012) |
| Mainland serow *Capricornis sumatraensis sumatraensis* | Thailand | Field survey | Relative abundance, species richness | DNP (2010) |
| | | Animal signs survey | MAXENT, | Trisurat et al. (2015); |
| | Malaysia | Camera trapping | Animal identification | Ain Ahmad Bakri et al. (2020) |
| *Capricornis sumatraensis milneedwardsii* | Thailand | Field survey | Relative abundance, species richness | DNP (2010) |
| | China | Animal signs survey | Habitat use | Chen et al. (2009) |

| **Species** | **Study area** | **Study methods** | **Habitat suitability analysis method** | **References** |
|---|---|---|---|---|
| | Vietnam | Transect survey, Interview | Mann – Whitney U tests | Thuc et al. (2014) |
| *Capricornis sumatraensis thar* | India | Camera trapping | Occupancy | Tapajit et al. (2012) |
| | Nepal | Animal signs | Population density, Ivelv's electivity index | Aryal (2009) |
| | | Animal signs, direct observation | Frequency of presence, intactness index | Paudel and Kindlmann (2012) |

Table S2 Number of species occurrences. The raw data and data after spatial thinning are shown by species and data collection method.

| | **Raw data** | | | | | **Thinned data** | | | | |
|---|---|---|---|---|---|---|---|---|---|---|
| Species | Camera trap | Direct Observation | Radio collar | Track/Sign | Total | Camera trap | Direct Observation | Radio collar | Track/Sign | Total |
| Gaur *Bos gaurus* | 3,055 | 286 | 0 | 22,436 | 25,777 | 257 | 247 | 0 | 2,228 | 2,732 |
| Banteng *Bos javanicus* | 483 | 16 | 4,341 | 911 | 5,751 | 64 | 16 | 34 | 247 | 361 |
| Wild water buffalo *Bubalus arnee* | 1 | 100 | 0 | 49 | 150 | 0 | 85 | 0 | 7 | 92 |
| Mainland serow *Capricornis sumatraensis* | 1,653 | 16 | 0 | 0 | 1,669 | 373 | 15 | 0 | 0 | 388 |
| Chinese goral *Naemorhedus griseus* | 291 | 26 | 0 | 0 | 3,17 | 125 | 10 | 0 | 0 | 135 |
| Total | 5,483 | 444 | 4,341 | 23,396 | 33,664 | 818 | 373 | 34 | 2,482 | 3,708 |

Table S3 Environmental factors use for modelling.

| No. | Variables | Description | Code | Source | Resolution | Spatial extent | Temporal range | CRS | Citation |
|---|---|---|---|---|---|---|---|---|---|
| 1 | Human population density | Total number of human population in grid cell (1 km) | hpop | WorldPop | ~1 $km^2$ | World | 2020 | WGS84 | Stevens et al. (2015) https://www.worldpop.org/geodata/summary?id=24777 |
| 2 | Slope | the rate of elevation change along the direction of the water flow, and calculated using a 3 × 3 cell moving window. | slope | OpenTopography | 250 $m^2$ | World | 2018 | WGS84 | Amatulli et al. (2020) http://spatial-ecology.net/dokuwiki/doku.php?id=topovar90m |
| 3 | Elevation (m) | World elevation | elev | Shuttle Radar Topography Mission (SRTM) | ~1 $km^2$ | World | 2000 | WGS84 | Fick & Hijmans (2017) https://www.worldclim.org/data/worldclim21.html# |
| 4 | Annual Mean Temperature | Bioclimatic variables | bio1 | WorldClim | ~1 $km^2$ | World | 1970-2000 | WGS84 | Fick & Hijmans (2017) https://www.worldclim.org/data/worldclim21.html# |
| 5 | Mean Diurnal Range (Mean of monthly (max temp - min temp)) | | bio2 | | | | | | |
| 6 | Isothermality (BIO2/BIO7) (×100) | | bio3 | | | | | | |
| 7 | Temperature Seasonality (standard deviation ×100) | | bio4 | | | | | | |
| 8 | Max Temperature of Warmest Month | | bio5 | | | | | | |
| 9 | Min Temperature of Coldest Month | | bio6 | | | | | | |
| 10 | Temperature Annual Range (BIO5-BIO6) | | bio7 | | | | | | |
| 11 | Mean Temperature of Wettest Quarter | | bio8 | | | | | | |
| 12 | Mean Temperature of Driest Quarter | | bio9 | | | | | | |

| No. | Variables | Description | Code | Source | Resolution | Spatial extent | Temporal range | CRS | Citation |
|---|---|---|---|---|---|---|---|---|---|
| 13 | Mean Temperature of Warmest Quarter | | bio10 | | | | | | |
| 14 | Mean Temperature of Coldest Quarter | | bio11 | | | | | | |
| 15 | Annual Precipitation | | bio12 | | | | | | |
| 16 | Annual Mean Temperature | | bio1 | | | | | | |
| 17 | Precipitation of Driest Month | | bio14 | | | | | | |
| 18 | Precipitation Seasonality (Coefficient of Variation) | | bio15 | | | | | | |
| 19 | Precipitation of Wettest Quarter | | bio16 | | | | | | |
| 20 | Precipitation of Driest Quarter | | bio17 | | | | | | |
| 21 | Precipitation of Warmest Quarter | | bio18 | | | | | | |
| 22 | Precipitation of Coldest Quarter | | bio19 | | | | | | |
| 23 | Normalized Difference Vegetation Index (NDVI) | NDVI (MODIS/Terra Vegetation Indices 16-Day L3 Global 1km-MODIS13A2), time series record for historical and climate applications | ndvi | MODIS, NASA | ~1 $km^2$ | Asia | 31-Dec-2019, time-series (16 days data) | sinusoidal | Didan, K. (2015) https://search.earthdata.nasa.gov/search?q=C194001238-LPDAAC_ECS |
| 24 | Crop | Crop cover fraction (%) | crop | Copernicus Global Land Service (CGLS): Land Cover 100m: epoch 2015: Globe, version2.0.2 | 100 m2 | World | 2015 | WGS84 | Buchhorn et al. (2019) https://zenodo.org/record/3243509#.YSve0C0RpQI |
| 25 | Grass | Grass cover fraction (%) | grass | | | World | 2015 | | |
| 26 | Tree | Tree cover fraction (%) | tree | | | World | 2015 | | |
| 27 | Urban | Urban cover fraction (%) | urban | | | World | 2015 | | |
| 28 | Water | Water cover fraction (%) | water | | | World | 2015 | | |

Table S4 Terrestrial ecoregions name by accessible areas using for modelling. The numbers mean that those accessible areas were included for modelling.

| | Accessible areas | | | | | | |
|---|---|---|---|---|---|---|---|
| | | species-specific accessible area | | | | | |
| Ecoregions | Large accessible areas | banteng | buffalo | gaur | goral | serow | Total |
| Afghan Mountains semi-desert | 1 | | | | | | 1 |
| Alashan Plateau semi-desert | 1 | | | | | | 1 |
| Altai alpine meadow and tundra | 1 | | | | | | 1 |
| Altai montane forest and forest steppe | 1 | | | | | | 1 |
| Altai steppe and semi-desert | 1 | | | | | | 1 |
| Amur meadow steppe | 1 | | | | | | 1 |
| Baluchistan xeric woodlands | 1 | | 1 | | | | 2 |
| Bohai Sea saline meadow | 1 | | | | | | 1 |
| Borneo lowland rain forests | 1 | 1 | | | | | 2 |
| Borneo montane rain forests | 1 | 1 | | | | | 2 |
| Borneo peat swamp forests | 1 | 1 | | | | | 2 |
| Brahmaputra Valley semi-evergreen forests | 1 | | 1 | 1 | 1 | 1 | 5 |
| Cardamom Mountains rain forests | 1 | | 1 | 1 | | 1 | 4 |
| Central Afghan Mountains xeric woodlands | | | 1 | | | | 1 |
| Central China loess plateau mixed forests | 1 | | | | 1 | | 2 |
| Central Deccan Plateau dry deciduous forests | 1 | | 1 | 1 | | | 3 |
| Central Indochina dry forests | 1 | 1 | 1 | 1 | 1 | 1 | 6 |
| Central Tibetan Plateau alpine steppe | 1 | | | | | | 1 |
| Changbai Mountains mixed forests | 1 | | | | | | 1 |
| Changjiang Plain evergreen forests | 1 | | | | 1 | 1 | 3 |
| Chao Phraya freshwater swamp forests | 1 | 1 | 1 | 1 | | 1 | 5 |
| Chao Phraya lowland moist deciduous forests | 1 | 1 | 1 | 1 | | 1 | 5 |
| Chhota-Nagpur dry deciduous forests | 1 | | 1 | 1 | | | 3 |
| Chin Hills-Arakan Yoma montane forests | 1 | | 1 | 1 | 1 | 1 | 5 |
| Da Hinggan-Dzhagdy Mountains conifer forests | 1 | | | | | | 1 |
| Daba Mountains evergreen forests | 1 | | | | 1 | 1 | 3 |
| Deccan thorn scrub forests | 1 | | 1 | 1 | | | 3 |
| East Afghan montane conifer forests | 1 | | 1 | | | | 2 |
| East Deccan dry-evergreen forests | 1 | | 1 | 1 | | | 3 |
| Eastern Gobi desert steppe | 1 | | | | | | 1 |
| Eastern highlands moist deciduous forests | 1 | | 1 | 1 | | | 3 |
| Eastern Himalayan alpine shrub and meadows | 1 | | 1 | 1 | 1 | 1 | 5 |
| Eastern Himalayan broadleaf forests | 1 | | 1 | 1 | 1 | 1 | 5 |
| Eastern Himalayan subalpine conifer forests | 1 | | 1 | 1 | 1 | 1 | 5 |
| Eastern Java-Bali montane rain forests | 1 | 1 | | | | | 2 |
| Eastern Java-Bali rain forests | 1 | | | | | | 1 |
| Emin Valley steppe | 1 | | | | | | 1 |
| Ghorat-Hazarajat alpine meadow | 1 | | | | | | 1 |
| Goadavari-Krishna mangroves | 1 | | 1 | 1 | | | 3 |
| Guizhou Plateau broadleaf and mixed forests | 1 | | | | 1 | 1 | 3 |
| Hainan Island monsoon rain forests | 1 | | | | 1 | 1 | 3 |
| Helanshan montane conifer forests | 1 | | | | | | 1 |
| Hengduan Mountains subalpine conifer forests | 1 | | | | 1 | 1 | 3 |

| Ecoregions | Accessible areas | | | | | | |
|---|---|---|---|---|---|---|---|
| | | species-specific accessible area | | | | | |
| | Large accessible areas | banteng | buffalo | gaur | goral | serow | Total |
| Himalayan subtropical broadleaf forests | 1 | | 1 | 1 | 1 | 1 | 5 |
| Himalayan subtropical pine forests | 1 | | 1 | 1 | | 1 | 4 |
| Hindu Kush alpine meadow | 1 | | | | | | 1 |
| Huang He Plain mixed forests | 1 | | | | 1 | | 2 |
| Indochina mangroves | 1 | 1 | 1 | 1 | | 1 | 5 |
| Indus River Delta-Arabian Sea mangroves | 1 | | 1 | 1 | | | 3 |
| Indus Valley desert | 1 | | 1 | | | | 2 |
| Irrawaddy dry forests | 1 | 1 | 1 | 1 | 1 | 1 | 6 |
| Irrawaddy freshwater swamp forests | 1 | 1 | 1 | 1 | 1 | 1 | 6 |
| Irrawaddy moist deciduous forests | 1 | 1 | 1 | 1 | 1 | 1 | 6 |
| Jian Nan subtropical evergreen forests | 1 | | | | 1 | 1 | 3 |
| Junggar Basin semi-desert | 1 | | | | | | 1 |
| Karakoram-West Tibetan Plateau alpine steppe | 1 | | 1 | | | 1 | 3 |
| Kayah-Karen montane rain forests | 1 | | 1 | 1 | 1 | 1 | 5 |
| Khathiar-Gir dry deciduous forests | 1 | | 1 | 1 | | | 3 |
| Kinabalu montane alpine meadows | 1 | 1 | | | | | 2 |
| Lower Gangetic Plains moist deciduous forests | 1 | | 1 | 1 | 1 | 1 | 5 |
| Luang Prabang montane rain forests | 1 | | 1 | 1 | 1 | 1 | 5 |
| Malabar Coast moist forests | 1 | | 1 | 1 | | | 3 |
| Manchurian mixed forests | 1 | | | | | | 1 |
| Meghalaya subtropical forests | 1 | | 1 | 1 | 1 | 1 | 5 |
| Mizoram-Manipur-Kachin rain forests | 1 | | 1 | 1 | 1 | 1 | 5 |
| Mongolian-Manchurian grassland | 1 | | | | | | 1 |
| Myanmar Coast mangroves | 1 | 1 | 1 | 1 | 1 | 1 | 6 |
| Myanmar coastal rain forests | 1 | | 1 | 1 | 1 | 1 | 5 |
| Narmada Valley dry deciduous forests | 1 | | 1 | 1 | | | 3 |
| Nenjiang River grassland | 1 | | | | | | 1 |
| North Tibetan Plateau-Kunlun Mountains alpine desert | 1 | | | | | | 1 |
| North Western Ghats moist deciduous forests | 1 | | 1 | 1 | | | 3 |
| North Western Ghats montane rain forests | 1 | | 1 | 1 | | | 3 |
| Northeast China Plain deciduous forests | 1 | | | | 1 | | 2 |
| Northeast India-Myanmar pine forests | 1 | | 1 | 1 | 1 | 1 | 5 |
| Northeastern Himalayan subalpine conifer forests | 1 | | 1 | 1 | 1 | 1 | 5 |
| Northern Annamites rain forests | 1 | | 1 | 1 | | 1 | 4 |
| Northern dry deciduous forests | 1 | | 1 | 1 | | | 3 |
| Northern Indochina subtropical forests | 1 | | 1 | 1 | 1 | 1 | 5 |
| Northern Khorat Plateau moist deciduous forests | 1 | | 1 | 1 | | 1 | 4 |
| Northern Thailand-Laos moist deciduous forests | 1 | | 1 | 1 | 1 | 1 | 5 |
| Northern Triangle subtropical forests | 1 | | 1 | 1 | 1 | 1 | 5 |
| Northern Triangle temperate forests | 1 | | 1 | 1 | 1 | 1 | 5 |
| Northern Vietnam lowland rain forests | 1 | | 1 | 1 | | 1 | 4 |
| Northwestern Himalayan alpine shrub and meadows | 1 | | 1 | | | 1 | 3 |
| Northwestern thorn scrub forests | 1 | | 1 | | | | 2 |
| Nujiang Langcang Gorge alpine conifer and mixed forests | 1 | | 1 | 1 | 1 | 1 | 5 |
| Ordos Plateau steppe | 1 | | | | 1 | | 2 |
| Orissa semi-evergreen forests | 1 | | 1 | 1 | | | 3 |

| | Accessible areas | | | | | | |
|---|---|---|---|---|---|---|---|
| | | species-specific accessible area | | | | | |
| Ecoregions | Large accessible areas | banteng | buffalo | gaur | goral | serow | Total |
| Pamir alpine desert and tundra | 1 | | | | | | 1 |
| Peninsular Malaysian montane rain forests | 1 | 1 | 1 | 1 | | 1 | 5 |
| Peninsular Malaysian peat swamp forests | 1 | 1 | 1 | 1 | | 1 | 5 |
| Peninsular Malaysian rain forests | 1 | | 1 | 1 | | 1 | 4 |
| Qaidam Basin semi-desert | 1 | | | | | | 1 |
| Qilian Mountains conifer forests | 1 | | | | 1 | 1 | 3 |
| Qilian Mountains subalpine meadows | 1 | | | | | | 1 |
| Qin Ling Mountains deciduous forests | 1 | | | | 1 | 1 | 3 |
| Qionglai-Minshan conifer forests | 1 | | | | 1 | 1 | 3 |
| Rann of Kutch seasonal salt marsh | 1 | | 1 | | | | 2 |
| Red River freshwater swamp forests | 1 | | 1 | | | 1 | 3 |
| Registan-North Pakistan sandy desert | | | 1 | | | | 1 |
| Rock and Ice | 1 | | 1 | 1 | 1 | 1 | 5 |
| Sichuan Basin evergreen broadleaf forests | 1 | | | | 1 | 1 | 3 |
| South China-Vietnam subtropical evergreen forests | 1 | | | | 1 | 1 | 3 |
| South Deccan Plateau dry deciduous forests | 1 | | 1 | 1 | | | 3 |
| South Iran Nubo-Sindian desert and semi-desert | 1 | | 1 | | | | 2 |
| South Western Ghats moist deciduous forests | 1 | | 1 | 1 | | | 3 |
| South Western Ghats montane rain forests | 1 | | 1 | 1 | | | 3 |
| Southeast Tibet shrublands and meadows | 1 | | | | 1 | 1 | 3 |
| Southeastern Indochina dry evergreen forests | 1 | | 1 | 1 | | 1 | 4 |
| Southern Annamites montane rain forests | 1 | 1 | 1 | 1 | | 1 | 5 |
| Southern Vietnam lowland dry forests | 1 | 1 | 1 | 1 | | 1 | 5 |
| Southwest Borneo freshwater swamp forests | 1 | 1 | | | | | 2 |
| Sri Lanka dry-zone dry evergreen forests | 1 | | 1 | | | | 2 |
| Sri Lanka lowland rain forests | 1 | | 1 | | | | 2 |
| Sri Lanka montane rain forests | 1 | | 1 | | | | 2 |
| Suiphun-Khanka meadows and forest meadows | 1 | | | | | | 1 |
| Sulaiman Range alpine meadows | 1 | | 1 | | | | 2 |
| Sumatran freshwater swamp forests | 1 | | | | | 1 | 2 |
| Sumatran lowland rain forests | 1 | | | | | 1 | 2 |
| Sumatran montane rain forests | 1 | | | | | 1 | 2 |
| Sumatran peat swamp forests | 1 | | | | | 1 | 2 |
| Sumatran tropical pine forests | 1 | | | | | 1 | 2 |
| Sunda Shelf mangroves | 1 | 1 | | | | 1 | 3 |
| Sundaland heath forests | 1 | 1 | | | | | 2 |
| Sundarbans freshwater swamp forests | 1 | | 1 | 1 | | | 3 |
| Sundarbans mangroves | 1 | | 1 | 1 | | | 3 |
| Taklimakan desert | 1 | | | | | | 1 |
| Tarim Basin deciduous forests and steppe | 1 | | | | | | 1 |
| Tenasserim-South Thailand semi-evergreen rain forests | 1 | 1 | 1 | 1 | | 1 | 5 |
| Terai-Duar savanna and grasslands | 1 | | 1 | 1 | 1 | 1 | 5 |
| Thar desert | 1 | | 1 | | | | 2 |
| Tian Shan montane conifer forests | 1 | | | | | | 1 |
| Tian Shan montane steppe and meadows | 1 | | | | | | 1 |
| Tibetan Plateau alpine shrublands and meadows | 1 | | | | | 1 | 2 |

| | Accessible areas | | | | | | |
|---|---|---|---|---|---|---|---|
| | | species-specific accessible area | | | | | |
| Ecoregions | Large accessible areas | banteng | buffalo | gaur | goral | serow | Total |
| Tonle Sap freshwater swamp forests | 1 | 1 | 1 | 1 | | 1 | 5 |
| Tonle Sap-Mekong peat swamp forests | 1 | 1 | 1 | 1 | | 1 | 5 |
| Upper Gangetic Plains moist deciduous forests | 1 | | 1 | 1 | | 1 | 4 |
| Western Himalayan alpine shrub and Meadows | 1 | | 1 | | | 1 | 3 |
| Western Himalayan broadleaf forests | 1 | | 1 | | | 1 | 3 |
| Western Himalayan subalpine conifer forests | 1 | | 1 | | | 1 | 3 |
| Western Java montane rain forests | 1 | 1 | | | | | 2 |
| Western Java rain forests | 1 | | | | | | 1 |
| Yarlung Tsangpo arid steppe | 1 | | 1 | | | 1 | 3 |
| Yellow Sea saline meadow | 1 | | | | | | 1 |
| Yunnan Plateau subtropical evergreen forests | 1 | | | 1 | 1 | 1 | 4 |
| Grand Total | 144 | 24 | 83 | 64 | 43 | 71 | 429 |

Table S5 Algorithms using in the modelling process (adapted from Andrade et al., 2020).

| **Algorithm** | **Abbreviations** | **Data used to create models** | **strength** | **limitations** | **Reference** |
|---|---|---|---|---|---|
| Bioclim (Envelope Score) | BIO | Presences-only | consider only climatic variables, not complicated using percentile distribution for prediction, easy to understand | has limitation on interpretation a non-climatic variables which can cause model inaccuracies | Pearson and Dawson (2003) |
| Generalized Linear Models | GLM | Presences and pseudo-absences | strong statistical foundation, flexible for modelling ecological relationship other than Gaussian distributions | use only for linear regression, not effective when occurrence data is small | Guisan et al. (2002) |
| Generalized Additive Models | GAM | Presences and pseudo-absences | fit highly non-linear and non-monotonic and complicated functions | not effective when occurrence data is small | Guisan et al. (2002); Ishihama et al. (2019) |
| Support Vector Machine | SVM | Presences and pseudo-absences | machine learning algorithm, reduce overprediction, alleviate multicollinearity | need absence (or pseudo-absence data) | Drake et al. (2006); Farrell et al. (2019) |
| Random Forest | RDF | Presences and pseudo-absences | machine learning algorithm, reduce overprediction, alleviate multicollinearity | non-parametric | Evans et al. (2011);Farrell et al. (2019) |
| Maximum Likelihood | MLK | Presences and background points | can use presence-only data, can estimate probability of occurrence using conventional likelihood methods | simulation can be slow, may have bias in small sample size | Royle et al. (2012) |
| Bayesian Gaussian Process | GAU | Presences and pseudo-absences | create joint models, conditional modelling, flexible, dealing with complex models | non-parametric, using presence/absence data, need experienced | Golding and Purse (2016) |
| Maximum Entropy default (all features) | MXD | Presences and background points | can use presence-only data, high performance which can differentiate between unsuitable and suitable area, can work with small sample size | cannot model a species fundamental niche, less localize, overfitting, prediction is not directly related to the actual parameter of interest, cannot estimate the probability of occurrence | Phillips et al. (2006) |

Table S6 Loading factors of the first two principal components (PC), PC1 and PC2, of the predictive environmental variables used in ecological niche modelling. LA is the the entire accessible areas and SSA is the species-specific accessible areas. The high values were highlighted in bold face.

| | | LA | | SSA | | | | | | | | | |
|---|---|---|---|---|---|---|---|---|---|---|---|---|---|
| | | All species | | *B. gaurus* | | *B. javanicus* | | *B. arnee* | | *C. sumatraensis* | | *N. griseus* | |
| | | PC1 | PC2 | PC1 | PC2 | PC1 | PC2 | PC1 | PC2 | PC1 | PC2 | PC1 | PC2 |
| Cumulative proportion | | 45% | 16% | 32% | 24% | 34% | 27% | 29% | 28% | 46% | 14% | 42% | 19% |
| Environmental variables | | PC1 | PC2 | PC1 | PC2 | PC1 | PC2 | PC1 | PC2 | PC1 | PC2 | PC1 | PC2 |
| Annual Mean Temperature | bio01 | **0.25** | 0.18 | **0.29** | 0.17 | 0.24 | 0.23 | **0.34** | 0.02 | **0.27** | 0.12 | **0.28** | 0.03 |
| Mean Diurnal Range | bio02 | -0.19 | 0.15 | 0.15 | **-0.26** | -0.17 | 0.10 | 0.02 | **-0.31** | -0.17 | 0.05 | **0.24** | 0.20 |
| Isothermality | bio03 | 0.21 | -0.15 | -0.02 | **0.29** | **0.28** | -0.12 | 0.08 | 0.24 | 0.14 | -0.26 | **0.27** | -0.08 |
| Temperature Seasonality | bio04 | -0.23 | 0.10 | 0.01 | **-0.32** | **-0.29** | 0.04 | -0.10 | **-0.29** | -0.19 | 0.15 | 0.24 | -0.15 |
| Max Temperature of Warmest Month | bio05 | 0.17 | **0.35** | **0.32** | -0.02 | 0.06 | **0.32** | 0.30 | -0.15 | 0.23 | 0.24 | 0.20 | -0.17 |
| Min Temperature of Coldest Month | bio06 | **0.27** | 0.06 | 0.19 | **0.29** | **0.31** | 0.07 | 0.29 | 0.18 | **0.27** | 0.03 | 0.14 | 0.17 |
| Temperature Annual Range | bio07 | -0.24 | 0.14 | 0.11 | **-0.32** | -0.28 | 0.12 | -0.03 | **-0.32** | -0.22 | 0.20 | -0.11 | -0.12 |
| Mean Temperature of Wettest Quarter | bio08 | 0.19 | **0.29** | **0.29** | 0.10 | 0.15 | **0.27** | 0.30 | -0.02 | 0.25 | 0.20 | 0.20 | -0.19 |
| Mean Temperature of Driest Quarter | bio09 | **0.25** | 0.10 | 0.28 | 0.16 | **0.29** | 0.15 | **0.32** | 0.02 | **0.26** | 0.08 | 0.15 | 0.15 |
| Mean Temperature of Warmest Quarter | bio10 | 0.20 | **0.31** | **0.32** | 0.05 | 0.13 | **0.30** | **0.32** | -0.10 | 0.24 | 0.21 | 0.17 | -0.16 |
| Mean Temperature of Coldest Quarter | bio11 | **0.26** | 0.08 | 0.24 | 0.25 | **0.29** | 0.14 | **0.31** | 0.13 | **0.27** | 0.05 | 0.16 | 0.14 |
| Annual Precipitation | bio12 | **0.25** | -0.16 | -0.09 | 0.25 | 0.20 | -0.18 | 0.03 | **0.31** | 0.23 | -0.15 | -0.20 | -0.10 |
| Precipitation of Wettest Month | bio13 | 0.21 | -0.04 | -0.03 | 0.17 | 0.03 | -0.07 | 0.05 | 0.23 | 0.19 | 0.00 | 0.05 | **-0.34** |
| Precipitation of Driest Month | bio14 | 0.17 | -0.24 | -0.09 | 0.20 | 0.24 | -0.19 | -0.03 | 0.19 | 0.13 | **-0.31** | -0.16 | **0.31** |
| Precipitation Seasonality | bio15 | 0.08 | 0.25 | 0.17 | -0.18 | -0.25 | 0.15 | 0.13 | -0.17 | -0.08 | **0.26** | 0.22 | 0.20 |
| Precipitation of Wettest Quarter | bio16 | 0.22 | -0.06 | -0.04 | 0.17 | 0.03 | -0.08 | 0.05 | 0.24 | 0.19 | -0.02 | **0.28** | -0.04 |
| Precipitation of Driest Quarter | bio17 | 0.18 | -0.24 | -0.09 | 0.21 | 0.25 | -0.19 | -0.04 | 0.21 | 0.14 | **-0.31** | -0.21 | 0.24 |
| Precipitation of Warmest Quarter | bio18 | 0.17 | -0.16 | -0.17 | 0.09 | -0.01 | -0.23 | -0.07 | 0.21 | 0.12 | -0.08 | 0.23 | 0.17 |
| Precipitation of Coldest Quarter | bio19 | 0.17 | -0.21 | -0.03 | 0.19 | 0.26 | -0.14 | 0.00 | 0.16 | 0.14 | **-0.31** | **0.27** | -0.07 |
| Crop cover fraction | crop | 0.06 | **0.28** | 0.23 | -0.12 | -0.03 | **0.26** | 0.18 | -0.15 | 0.06 | **0.35** | 0.01 | **0.27** |
| Elevation | elev | -0.17 | -0.26 | -0.27 | -0.18 | -0.22 | -0.22 | **-0.33** | -0.04 | **-0.25** | -0.16 | -0.21 | -0.25 |
| Grass cover fraction | grass | -0.11 | -0.09 | -0.04 | -0.12 | -0.05 | 0.12 | -0.10 | -0.05 | -0.20 | -0.07 | -0.21 | -0.10 |
| Human population density | hlog | 0.16 | 0.11 | 0.23 | -0.07 | -0.03 | 0.21 | 0.21 | -0.08 | 0.15 | 0.22 | 0.12 | **0.26** |
| Normalized Difference Vegetation Index | ndvi | 0.23 | -0.02 | -0.15 | 0.19 | 0.01 | -0.26 | 0.05 | 0.27 | 0.21 | -0.10 | 0.22 | -0.17 |
| Slope | slope | -0.02 | **-0.29** | **-0.28** | -0.03 | -0.14 | -0.23 | -0.26 | 0.11 | -0.12 | -0.20 | -0.06 | **-0.27** |
| Tree cover fraction | tree | 0.17 | -0.19 | -0.21 | 0.18 | 0.04 | **-0.29** | -0.06 | **0.27** | 0.15 | -0.23 | 0.16 | -0.21 |
| Urban cover fraction | urban | 0.04 | 0.08 | 0.05 | 0.02 | 0.02 | 0.10 | 0.05 | 0.00 | 0.03 | 0.12 | 0.02 | 0.20 |
| Water cover fraction | water | 0.00 | 0.01 | 0.01 | 0.02 | 0.02 | 0.04 | 0.01 | 0.01 | 0.01 | 0.03 | 0.02 | 0.05 |

Table S7 True Skill Statistics (TSS) and Area Under the Curve (AUC) values of the weighted average ensemble, and the threshold values for binary maps for five species classified by accessible area type and MSDM method.

Best performing models for each accessible area by TSS are shown in **Boldface**.

| | Large accessible area | | | | | | Species specific accessible area | | | | | |
|---|---|---|---|---|---|---|---|---|---|---|---|---|
| | No MSDMa | | | MSDM (OBR)b | | | No MSDMa | | | MSDM (OBR)b | | |
| | TSS | | AUC | TSS | | AUC | TSS | | AUC | TSS | | AUC |
| Species | Score | Threshold | Score | Score | Threshold | Score | Score | Threshold | Score | Score | Threshold | Score |
| Gaur *Bos gaurus* | **0.92** | 0.49 | 0.99 | 0.92 | 0.44 | 0.99 | **0.88** | 0.39 | 0.98 | 0.88 | 0.41 | 0.98 |
| Banteng *Bos javanicus* | 0.93 | 0.55 | 0.99 | **0.94** | 0.41 | 1 | 0.85 | 0.33 | 0.96 | **0.83** | 0.42 | 0.97 |
| Wild water buffalo *Bubalus arnee* | 0.67 | 0.47 | 0.88 | **0.72** | 0.6 | 0.9 | 0.57 | 0.58 | 0.83 | **0.85** | 0.44 | 0.95 |
| Mainland serow *Capricornis sumatraensis* | **0.87** | 0.55 | 0.97 | 0.76 | 0.47 | 0.94 | **0.93** | 0.57 | 0.98 | 0.93 | 0.52 | 0.98 |
| Chinese goral *Naemorhedus griseus* | 0.91 | 0.29 | 0.98 | **0.91** | 0.59 | 0.98 | 0.87 | 0.47 | 0.96 | **0.9** | 0.39 | 0.97 |

### Table S8 Suitable areas calculated from the best model and classified by country.

| species | Country | Country area (sq.km) | Species-specific accessible areas: Suitable area per country (sq.km) | (%) | Large accessible areas: Suitable area per country (sq.km) | (%) |
|---|---|---|---|---|---|---|
| Gaur | Bangladesh | 147,570 | 0 | 0 | 180 | 0 |
| (*Bos gaurus*) | Bhutan | 38,394 | 0 | 0 | 117 | 0 |
| | Cambodia | 181,035 | 38,465 | 13 | 59,140 | 10 |
| | China | 9,706,961 | 0 | 0 | 1 | 0 |
| | India | 3,287,590 | 44,856 | 15 | 176,830 | 29 |
| | Indonesia | 1,904,569 | 0 | 0 | 14,090 | 2 |
| | Laos | 236,800 | 21,838 | 7 | 41,553 | 7 |
| | Malaysia | 329,847 | 5,285 | 2 | 3,389 | 1 |
| | Myanmar | 676,578 | 45,829 | 15 | 97,763 | 16 |
| | Nepal | 147,181 | 1 | 0 | 2,097 | 0 |
| | Sri Lanka | 65,610 | 0 | 0 | 12,475 | 2 |
| | Thailand | 513,120 | 138,892 | 46 | 177,216 | 29 |
| | Vietnam | 331,210 | 7,853 | 3 | 26,343 | 4 |
| | Total | | **303,019** | **100** | **611,192** | **100** |
| Banteng | Bangladesh | 147,570 | 35 | 0 | 0 | 0 |
| (*Bos javanicus*) | Brunei Darussalam | 5,765 | 0 | 0 | 0 | 0 |
| | Cambodia | 181,035 | 68,463 | 24 | 87,388 | 20 |
| | China | 9,706,961 | 20 | 0 | 5 | 0 |
| | India | 3,287,590 | 4 | 0 | 0 | 0 |
| | Indonesia | 1,904,569 | 5,209 | 2 | 10,801 | 2 |
| | Laos | 236,800 | 14,337 | 5 | 31,742 | 7 |
| | Malaysia | 329,847 | 5,985 | 2 | 4,652 | 1 |
| | Myanmar | 676,578 | 39,596 | 14 | 78,203 | 18 |
| | Thailand | 513,120 | 125,307 | 44 | 180,275 | 40 |
| | Vietnam | 331,210 | 23,761 | 8 | 52,150 | 12 |
| | Total | | **282,717** | **100** | **445,216** | **100** |
| Wild water buffalo | Bangladesh | 147,570 | 90,097 | 15 | 75,175 | 25 |
| (*Bubalus arnee*) | Bhutan | 38,394 | 846 | 0 | 324 | 0 |
| | Cambodia | 181,035 | 283 | 0 | 14 | 0 |
| | China | 9,706,961 | 546 | 0 | 106 | 0 |
| | India | 3,287,590 | 213,247 | 35 | 106,149 | 35 |
| | Laos | 236,800 | 16,353 | 3 | 244 | 0 |
| | Malaysia | 329,847 | 3,744 | 1 | 0 | 0 |
| | Myanmar | 676,578 | 99,844 | 16 | 63,246 | 21 |
| | Nepal | 147,181 | 11,729 | 2 | 5,642 | 2 |
| | Sri Lanka | 65,610 | 43,010 | 7 | 27,254 | 9 |
| | Thailand | 513,120 | 135,552 | 22 | 28,450 | 9 |
| | Total | | **615,250** | **100** | **306,603** | **100** |
| Mainland serow | Bangladesh | 147,570 | 9 | 0 | 6 | 0 |
| (*Capricornis sumatraensis*) | Bhutan | 5,765 | 1,167 | 0 | 1,178 | 0 |
| | Cambodia | 181,035 | 40,718 | 14 | 31,572 | 7 |
| | China | 9,706,961 | 55,528 | 19 | 60,890 | 13 |
| | India | 3,287,590 | 5,709 | 2 | 84,742 | 18 |
| | Indonesia | 1,904,569 | 3 | 0 | 1,259 | 0 |
| | Laos | 236,800 | 37,884 | 13 | 27,309 | 6 |
| | Malaysia | 329,847 | 7,318 | 2 | 11,170 | 2 |
| | Myanmar | 676,578 | 66,892 | 23 | 122,728 | 26 |
| | Nepal | 147,181 | 275 | 0 | 305 | 0 |
| | Sri Lanka | 65,610 | 0 | 0 | 9,895 | 2 |
| | Thailand | 513,120 | 68,534 | 23 | 109,782 | 23 |
| | Vietnam | 331,210 | 9,708 | 3 | 19,784 | 4 |
| | Total | | **293,746** | **100** | **480,620** | **100** |
| Chinese goral | China | 9,706,961 | 285,073 | 86 | 162,929 | 96 |
| (*Naemorhedus griseus*) | India | 3,287,590 | 15,413 | 5 | 1,067 | 1 |
| | Myanmar | 676,578 | 16,369 | 5 | 5,884 | 3 |
| | Thailand | 513,120 | 14,847 | 4 | 0 | 0 |
| | Total | | **331,701** | **100** | **169,881** | **100** |

Table S9 Habitat suitability areas classified by IUCN protected are categories 1 to 7, not applicable and non protected areas.

| | | Species-specific accessible areas | | | Large accessible areas | | |
|---|---|---|---|---|---|---|---|
| | | | Suitable areas overlap with IUCN category | | | Suitable areas overlap with IUCN category | |
| species | IUCN protected areas category | Model | (sq.km) | % | Model | (sq.km) | % |
| Gaur | IUCN PA Ia, Ib | No MSDM | 28,904 | 10 | No MSDM | 39,219 | 6 |
| *(Bos gaurus*) | IUCN PA II | | 50,497 | 17 | | 67,587 | 11 |
| | IUCN PA III | | 3,919 | 1 | | 291 | 0 |
| | IUCN PA IV | | 17,714 | 6 | | 31,345 | 5 |
| | IUCN PA V | | 492 | 0 | | 607 | 0 |
| | IUCN PA VI | | 14,689 | 5 | | 19,986 | 3 |
| | Not applicable | | 6,570 | 2 | | 7,057 | 1 |
| | Not protected area | | 180,380 | 60 | | 446,456 | 73 |
| | Total | | 303,165 | 100 | | 612,548 | 100 |
| Banteng | IUCN PA Ia, Ib | No MSDM | 21,987 | 8 | OBR | 30,500 | 7 |
| *(Bos javanicus*) | IUCN PA II | | 42,459 | 15 | | 57,894 | 13 |
| | IUCN PA III | | 4,141 | 1 | | 253 | 0 |
| | IUCN PA IV | | 30,649 | 11 | | 32,650 | 7 |
| | IUCN PA V | | 2,646 | 1 | | 520 | 0 |
| | IUCN PA VI | | 19,852 | 7 | | 21,879 | 5 |
| | Not applicable | | 6,272 | 2 | | 2,094 | 0 |
| | Not protected area | | 156,505 | 55 | | 300,285 | 67 |
| | Total | | 284,510 | 100 | | 446,075 | 100 |
| Wild water buffalo | IUCN PA Ia, Ib | OBR | 16,623 | 3 | OBR | 8,815 | 3 |
| *(Bubalus arnee*) | IUCN PA II | | 17,677 | 3 | | 8,551 | 3 |
| | IUCN PA III | | 0 | 0 | | 0 | 0 |
| | IUCN PA IV | | 12,521 | 2 | | 4,137 | 1 |
| | IUCN PA V | | 19 | 0 | | 8 | 0 |
| | IUCN PA VI | | 2,528 | 0 | | 505 | 0 |
| | Not applicable | | 6,010 | 1 | | 5,746 | 2 |
| | Not protected area | | 560,521 | 91 | | 279,180 | 91 |
| | Total | | 615,899 | 100 | | 306,942 | 100 |
| Mainland serow | IUCN PA Ia, Ib | No MSDM | 23,034 | 8 | No MSDM | 35,965 | 7 |
| *(Capricornis sumatraensis*) | IUCN PA II | | 44,754 | 15 | | 59,875 | 12 |
| | IUCN PA III | | 1,344 | 0 | | 349 | 0 |
| | IUCN PA IV | | 18,760 | 6 | | 16,894 | 4 |
| | IUCN PA V | | 2,683 | 1 | | 265 | 0 |
| | IUCN PA VI | | 19,960 | 7 | | 14,170 | 3 |
| | Not applicable | | 9,195 | 3 | | 11,995 | 2 |
| | Not protected area | | 174,153 | 59 | | 341,728 | 71 |
| | Total | | 293,883 | 100 | | 481,241 | 100 |
| Chinese goral | IUCN PA Ia, Ib | OBR | 3,368 | 1 | OBR | 0 | 0 |
| *(Naemorhedus griseus*) | IUCN PA II | | 6,635 | 2 | | 2,886 | 2 |
| | IUCN PA III | | 0 | 0 | | 0 | 0 |
| | IUCN PA IV | | 611 | 0 | | 33 | 0 |
| | IUCN PA V | | 0 | 0 | | 0 | 0 |
| | IUCN PA VI | | 1,250 | 0 | | 985 | 1 |
| | Not applicable | | 23,749 | 7 | | 20,332 | 12 |
| | Not protected area | | 296,091 | 89 | | 145,646 | 86 |
| | Total | | 331,704 | 100 | | 169,882 | 100 |

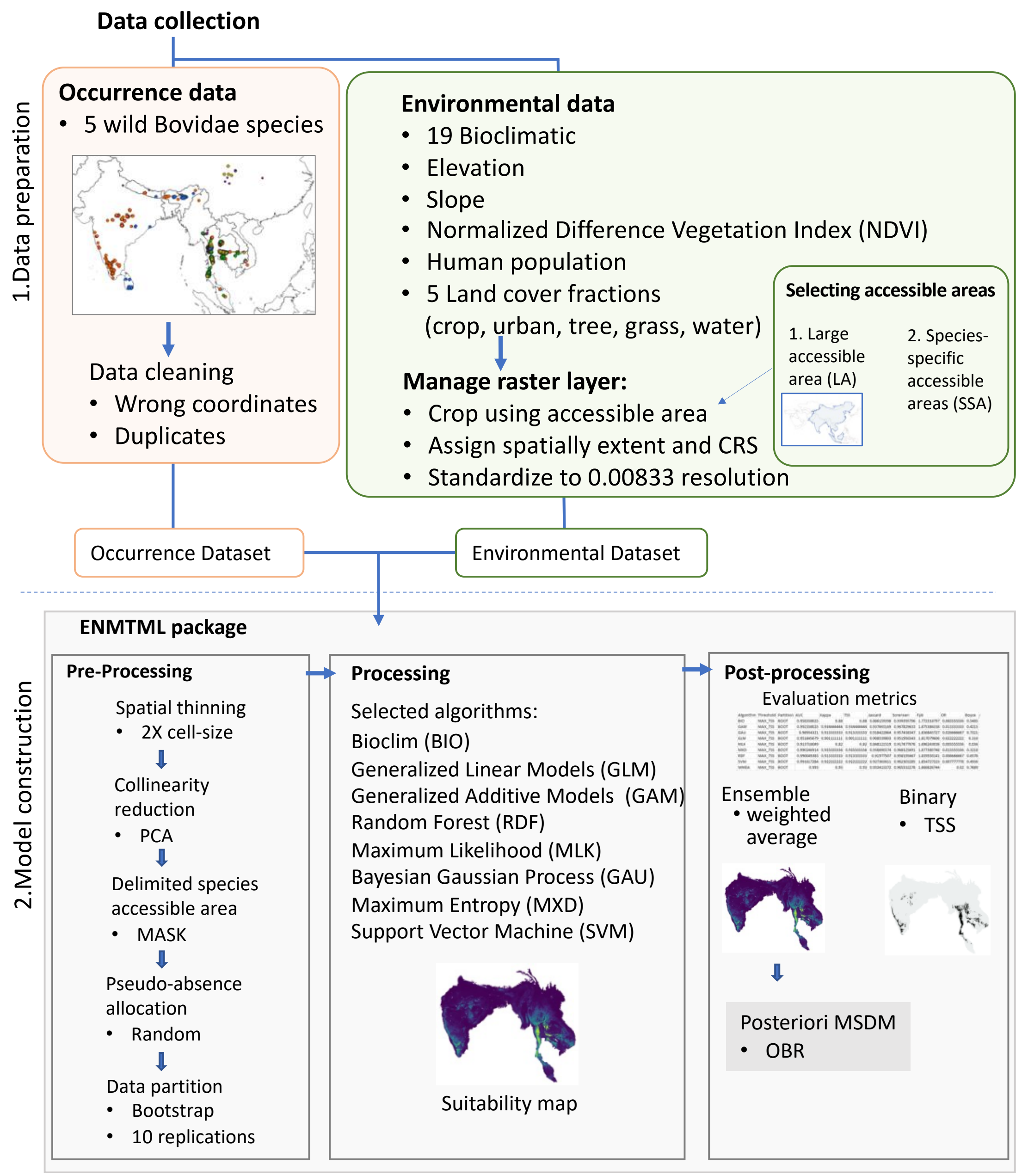


Figure S1. Our study workflow was based on the frameworks from Andrade et al. (2020) and Peterson et al. (2011).

Figure S2. Principal Component Analysis (PCA) for gaur (Bos Gaurus), banteng (Bos javanicus), wild water buffalo (Bubalus arnee), mainland serow(Capricornis sumatraensis) and Chinese goral (Naemorhedus griseus).

(A) The negative (red) and positive (blue) relationships between the loading of important principal components (PC) that are used in model building and environmental variables.

(B) The first two PCA axes (PC 1 and PC2) of the most explained variance plotted with variable loadings (arrows) and species' locations. The loadings arrows represent relationships and importance of environmental variables and species presence-absence (the longer line = more important with the negative-positive influence). The points represent species' locations for 4 categories: 00 (light blue) = absence observation, absence prediction; 01 (blue) = absence observation; 10 (purple) = presence observation, absence prediction, 11 (pink) = presence observation, presence prediction.

(C) Scree plot of explained variance for important PC (%).

1. Gaur (Bos gaurus)

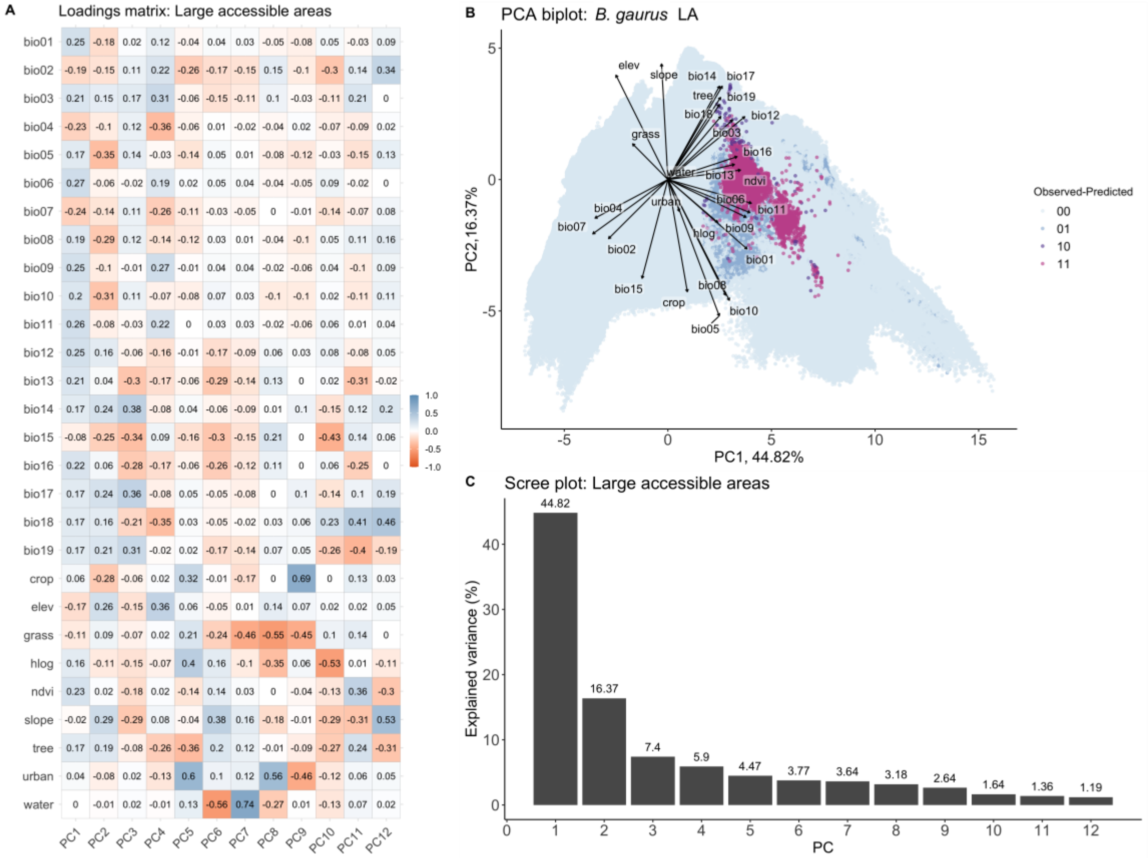

2. Banteng (*Bos javanicus*)

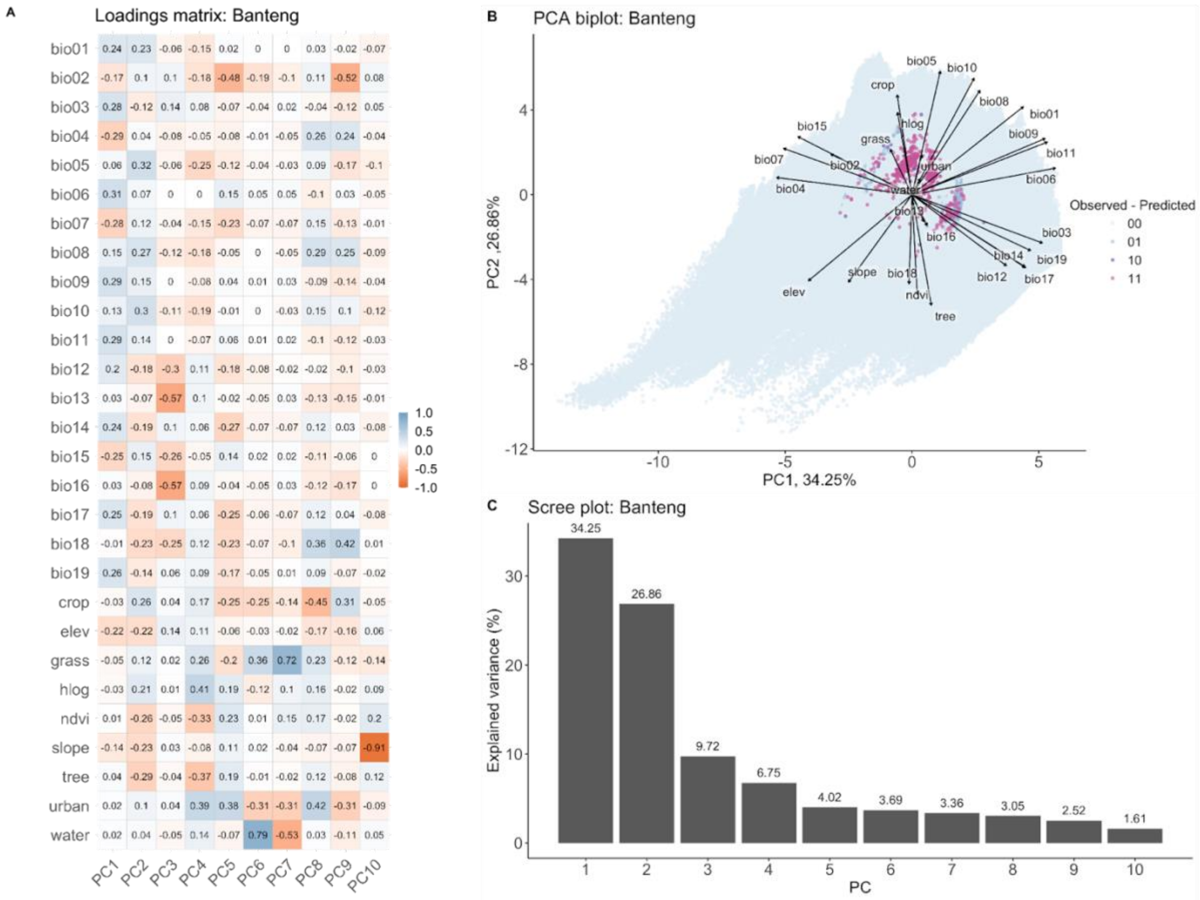

3. Wild water buffalo (*Bubalus arnee*)

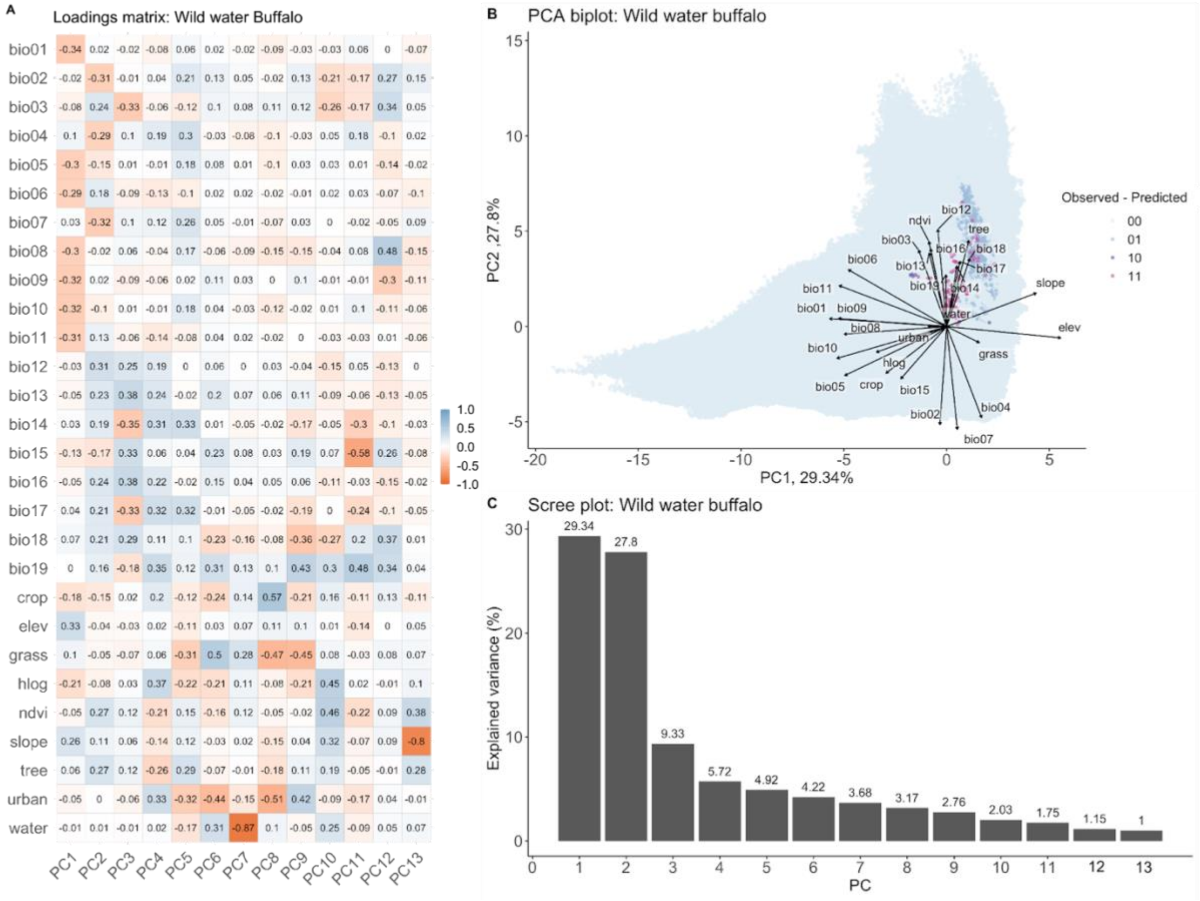

4. Mainland serow (Capricornis sumatraensis)

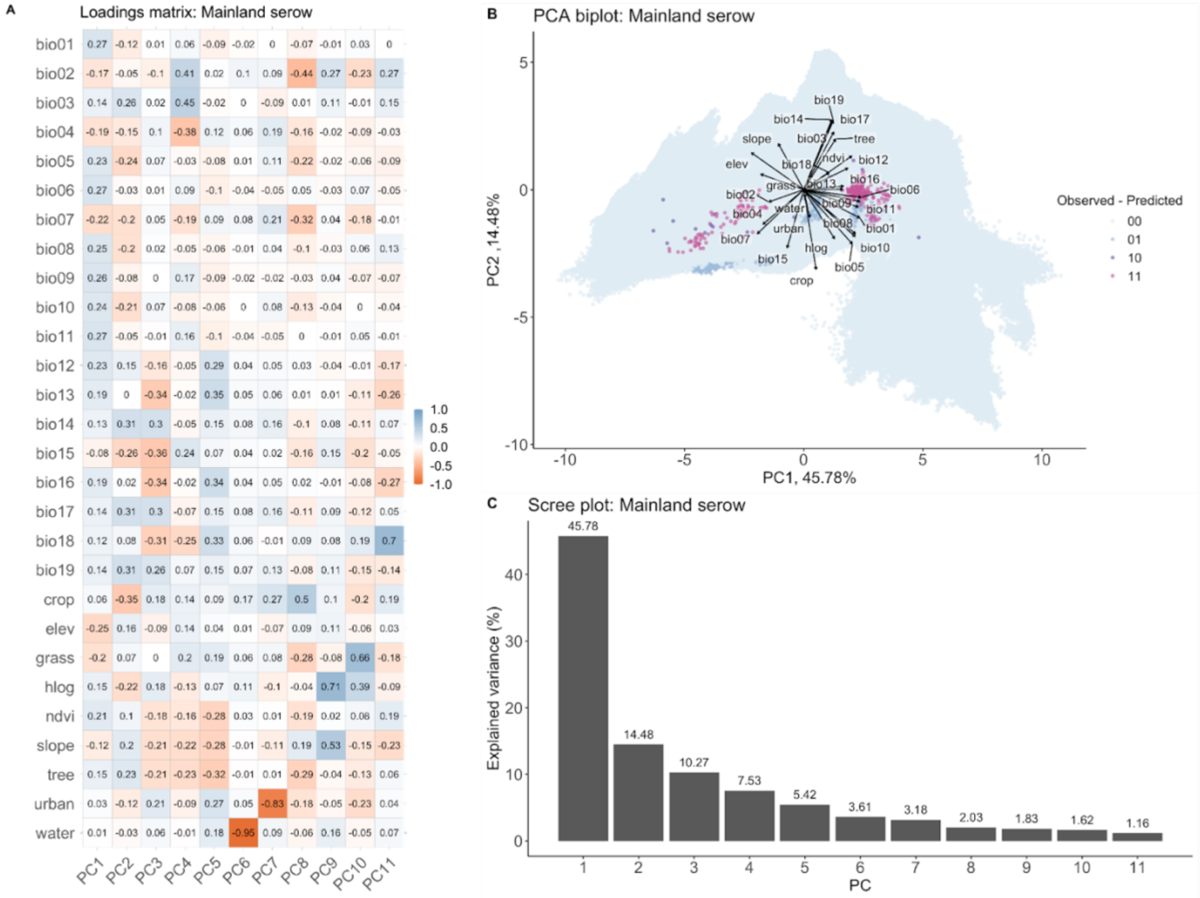

5. Chinese goral (*Naemorhedus griseus*)

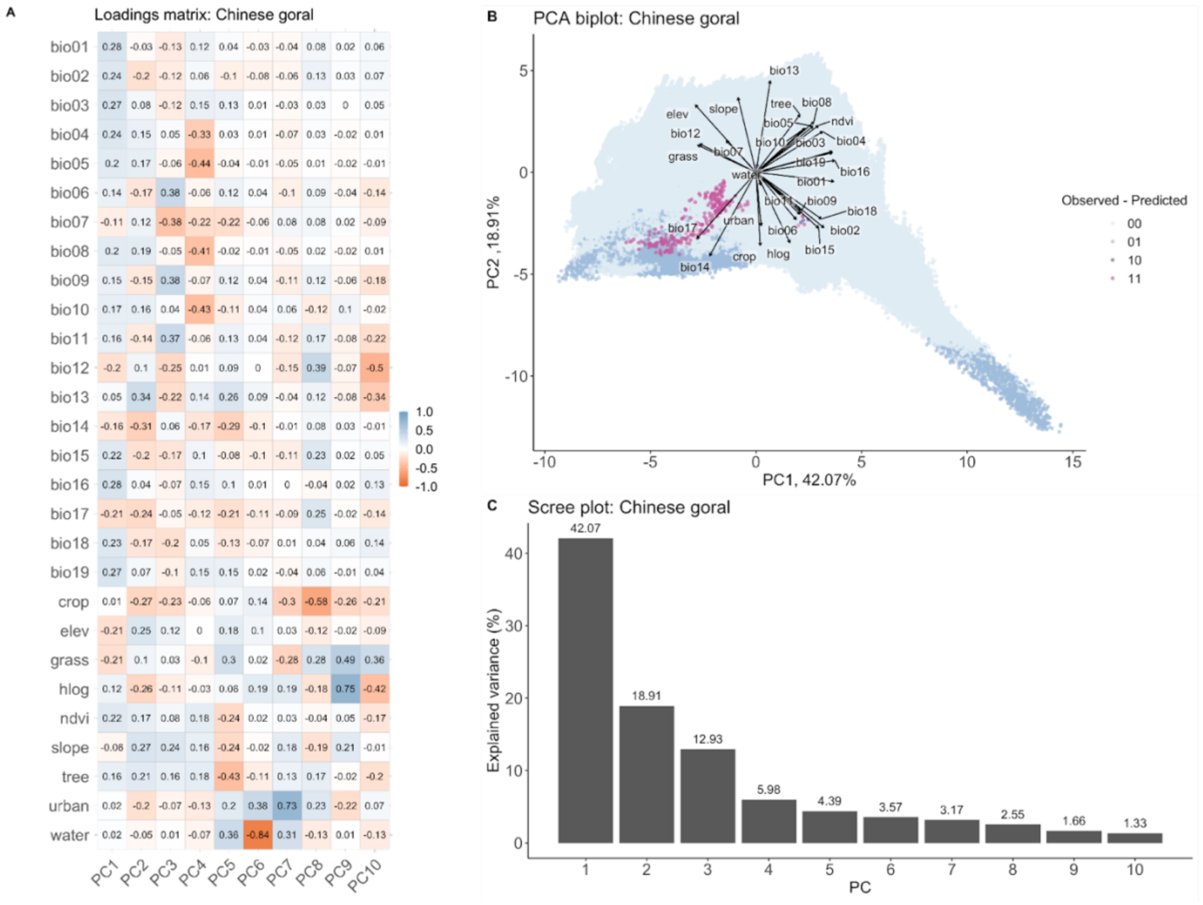

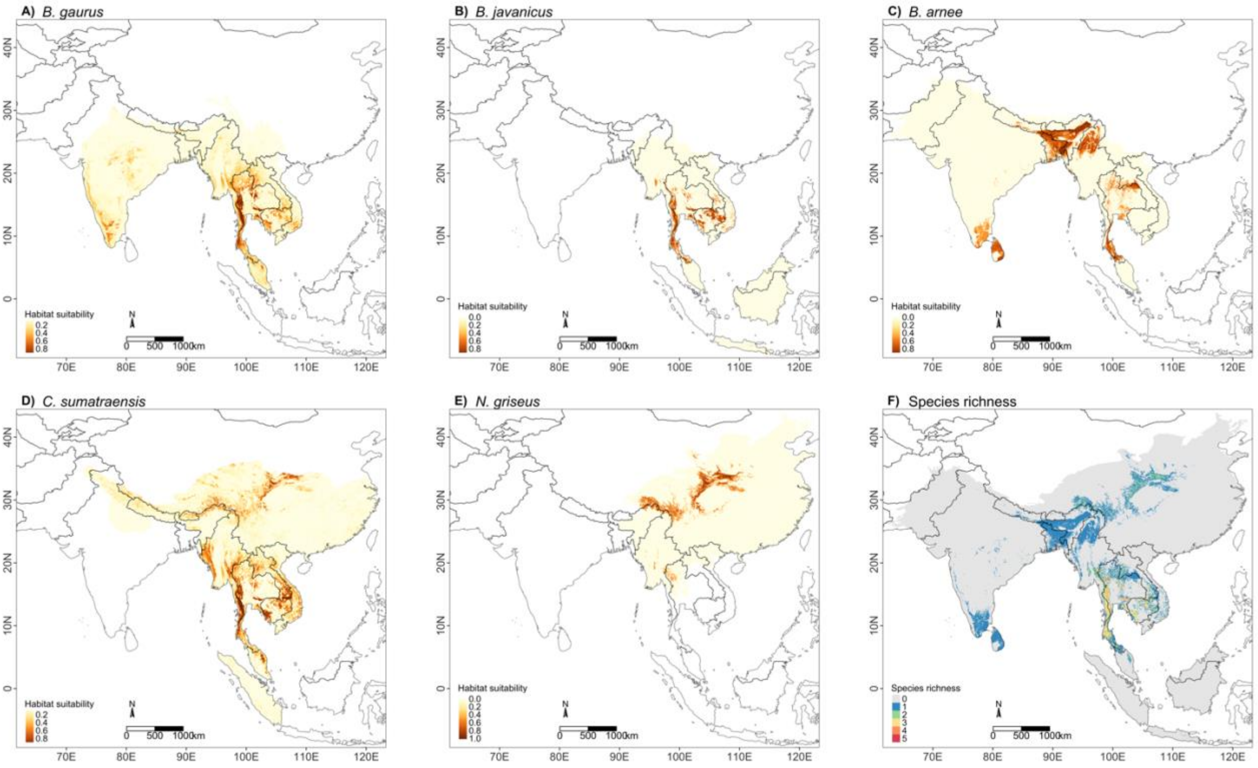


Figure S3. (Top) Habitat suitability prediction maps of the best-performing models for five species (A-E) using species-specific accessible areas and weighted average ensemble models. The value ranges from 0-1: yellow represents low suitability and dark brown represents high suitability. (F) presents the species richness map for five species using the binary suitability results, ranging from 0 (not suitable area for the five species) to 5 (suitable area for all five species). (Bottom) Comparison of no MSDM and OBR spatially restrict accessible areas of the suitable areas of SSA models classified by protected and not protected areas.

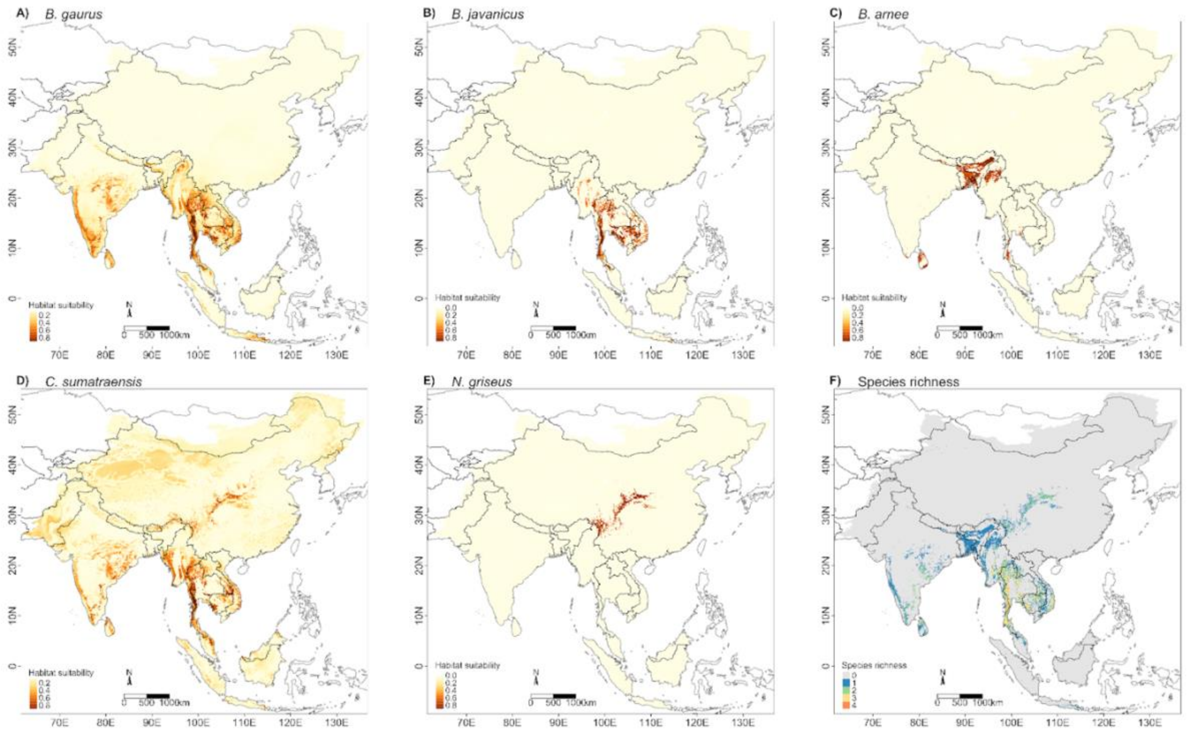


Figure S4. (Top) Habitat suitability prediction maps for five species (A-E) using the large accessible area. The value ranges from 0-1: yellow represents low suitability and dark brown represents high suitability. (F) Species richness map for five species which range from 0 (no suitable area for the five species) to 4 (suitable area for four species), which is the highest value range for the large accessible area model predictions. (Bottom) Comparison of no MSDM and OBR spatially restrict accessible areas of the suitable areas of LA models classified by protected and not protected areas.

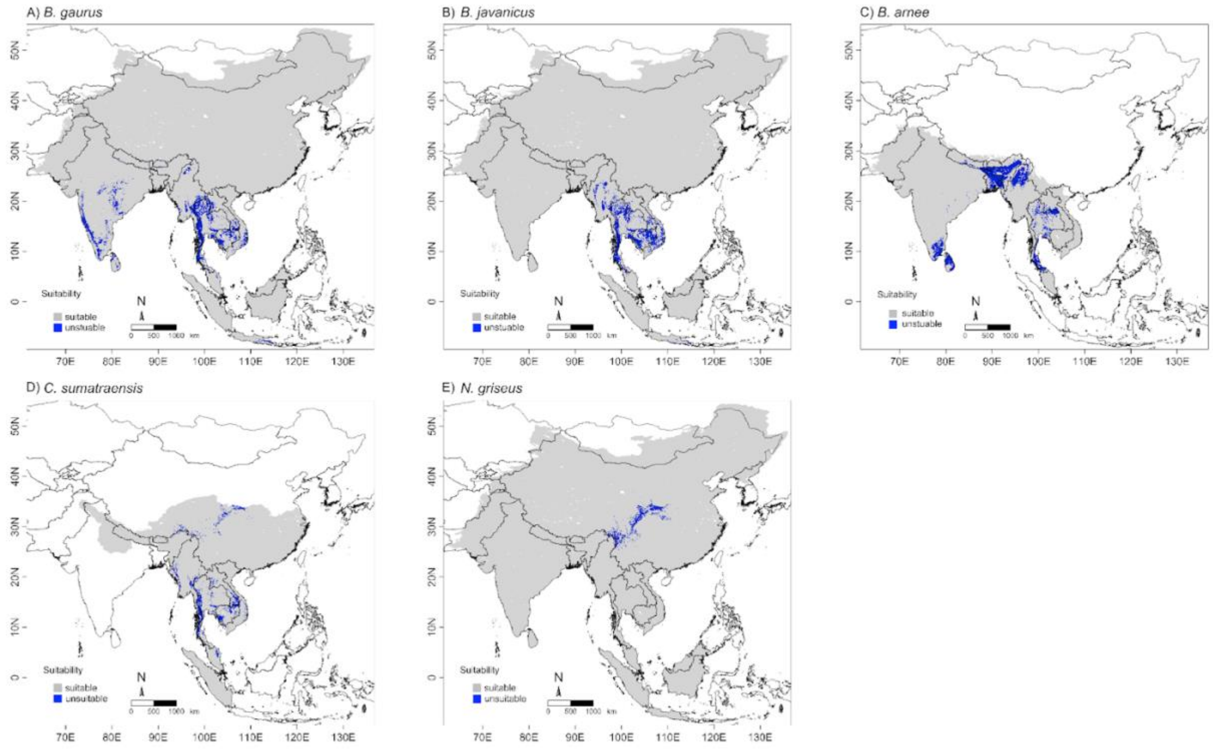


Figure S5. The best TSS models of binary maps of suitable areas for five species (A-E). The grey color is unsuitable and blue is suitable areas.

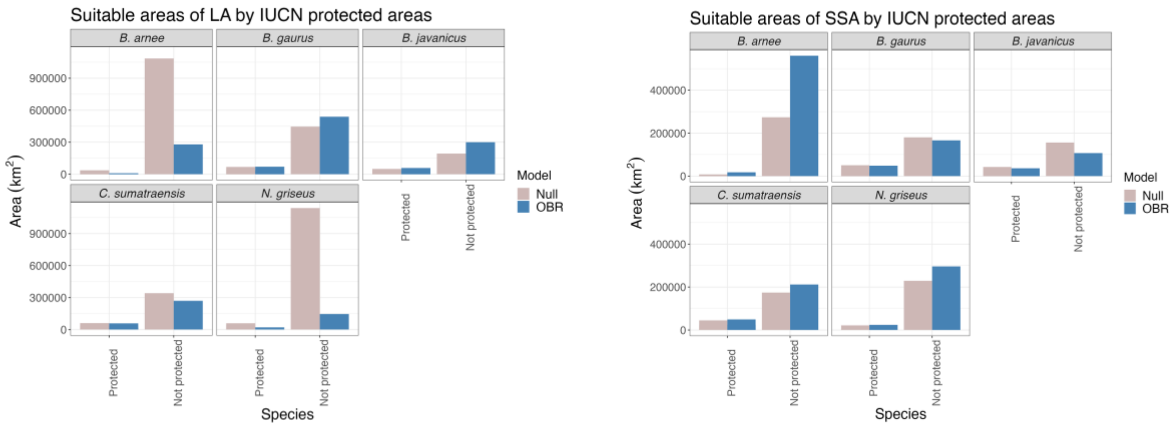


Figure S6. The suitable area in each species classified by IUCN protected areas (PA) and non PA between two types of accessible areas A) LA and B) SSA and spatially restrictions: Null (No MSDM; pink) and OBR (occurrences-based threshold (OBR; blue). All models have the suitable areas outside IUCN PA, and the largest one outside PAs is *B.arneee*. OBR has similar results with Null in *B. gaurs, B. javanicus and C. sumatraensis* but show a huge difference for B. arnee (both LA and SSA) and N. griseus (LA). Thus, we recommend using the SSA for small and confine species to increase the model precision.

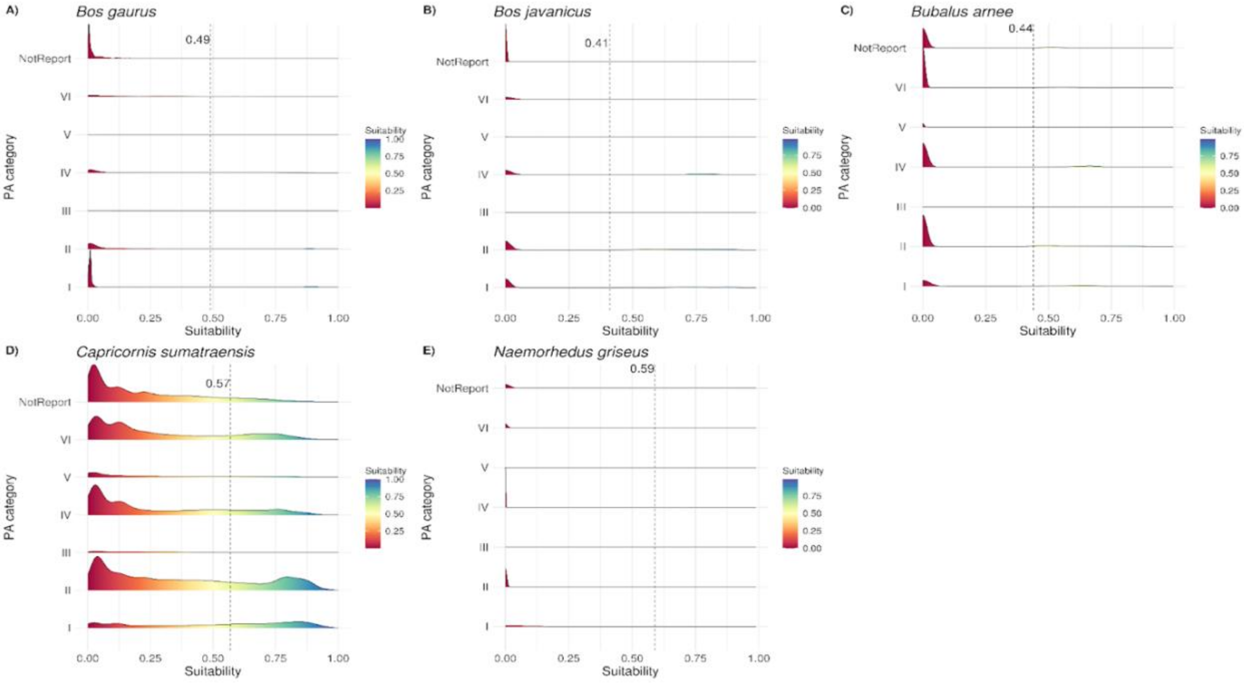


Figure S7. Density of suitability for five species (A - E) by IUCN protected areas category (1 to 6) and not report or not applicable protected areas. The values range from 0 (low suitability - blue) to 1 (high suitability - red). The dashed lines show the mean threshold values calculated for the best performing models for each species.

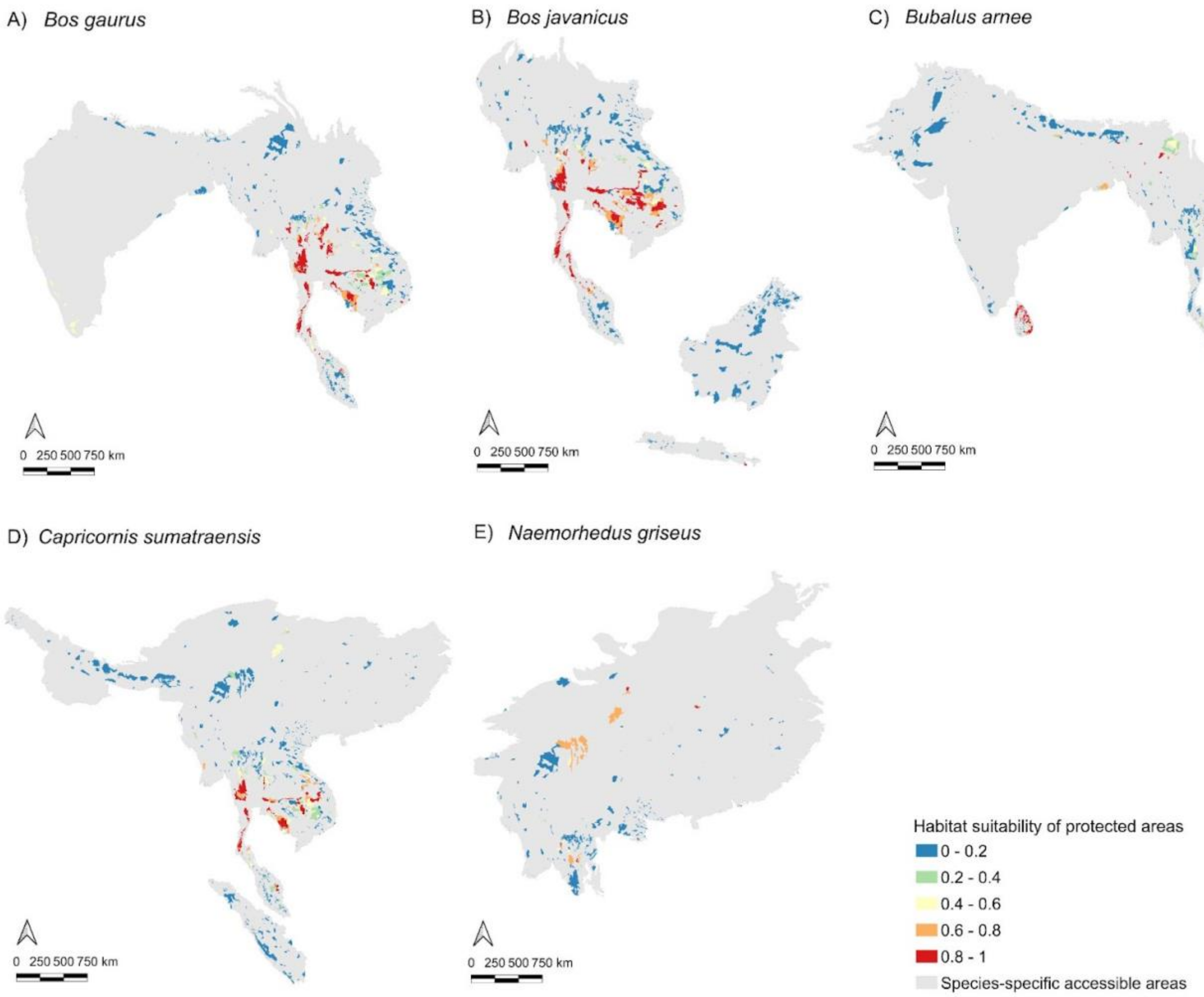


Figure S8.The proportion of suitable areas range from 0 (unsuitable) to 1 (high proportion of suitable) with species accessible area model for five species and their best performing models: (A) Guar (*Bos gaurus*), (B) Banteng (*Bos javanicus*), (C) wild water buffalo (*Bubalus arnee*), (D) mainland serow (*Capricornis sumatraensis*), and (E) Chinese goral (*Naemorhedus griseus*).

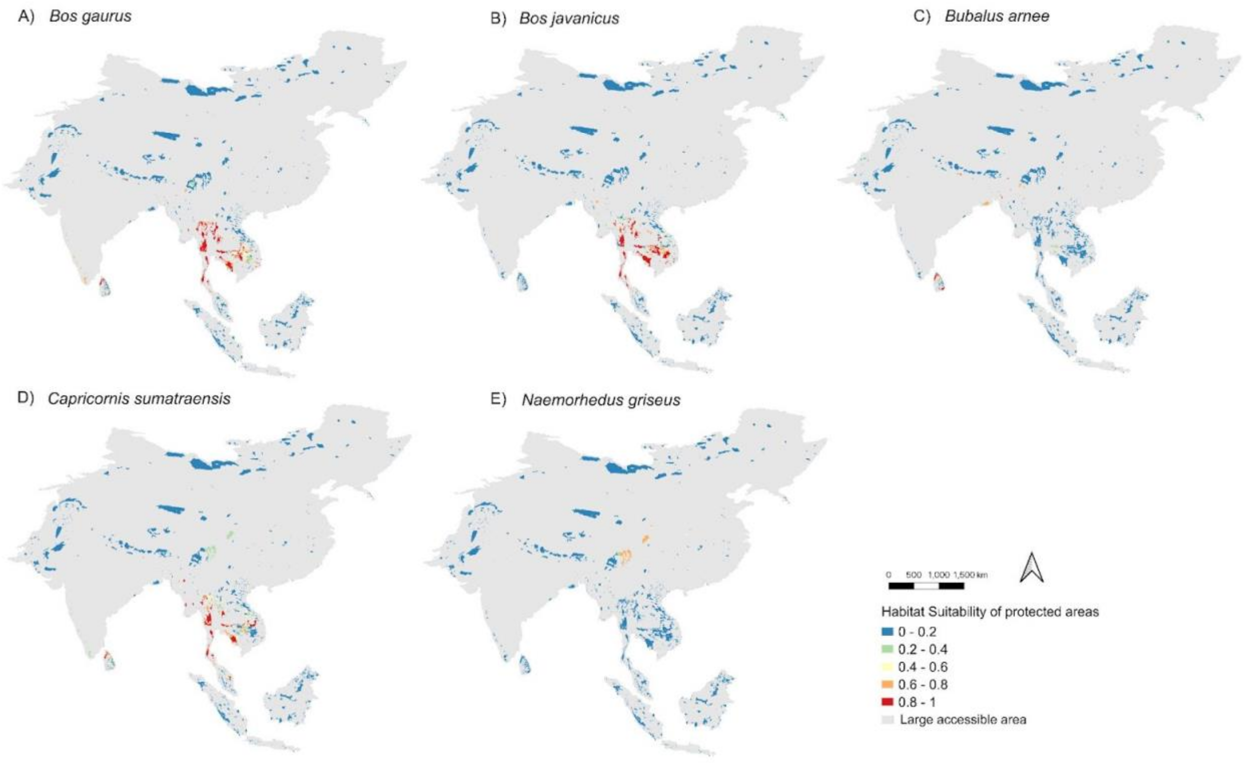


Figure S9 The proportion of suitable areas range from 0 (unsuitable) to 1 (high proportion of suitable) with large accessible area models for five species and their best performing models: (A) Guar (*Bos gaurus*), (B) Banteng (*Bos javanicus*), (C) wild water buffalo (*Bubalus arnee*), (D) mainland serow (*Capricornis sumatraensis*), and (E) Chinese goral (*Naemorhedus griseus*).

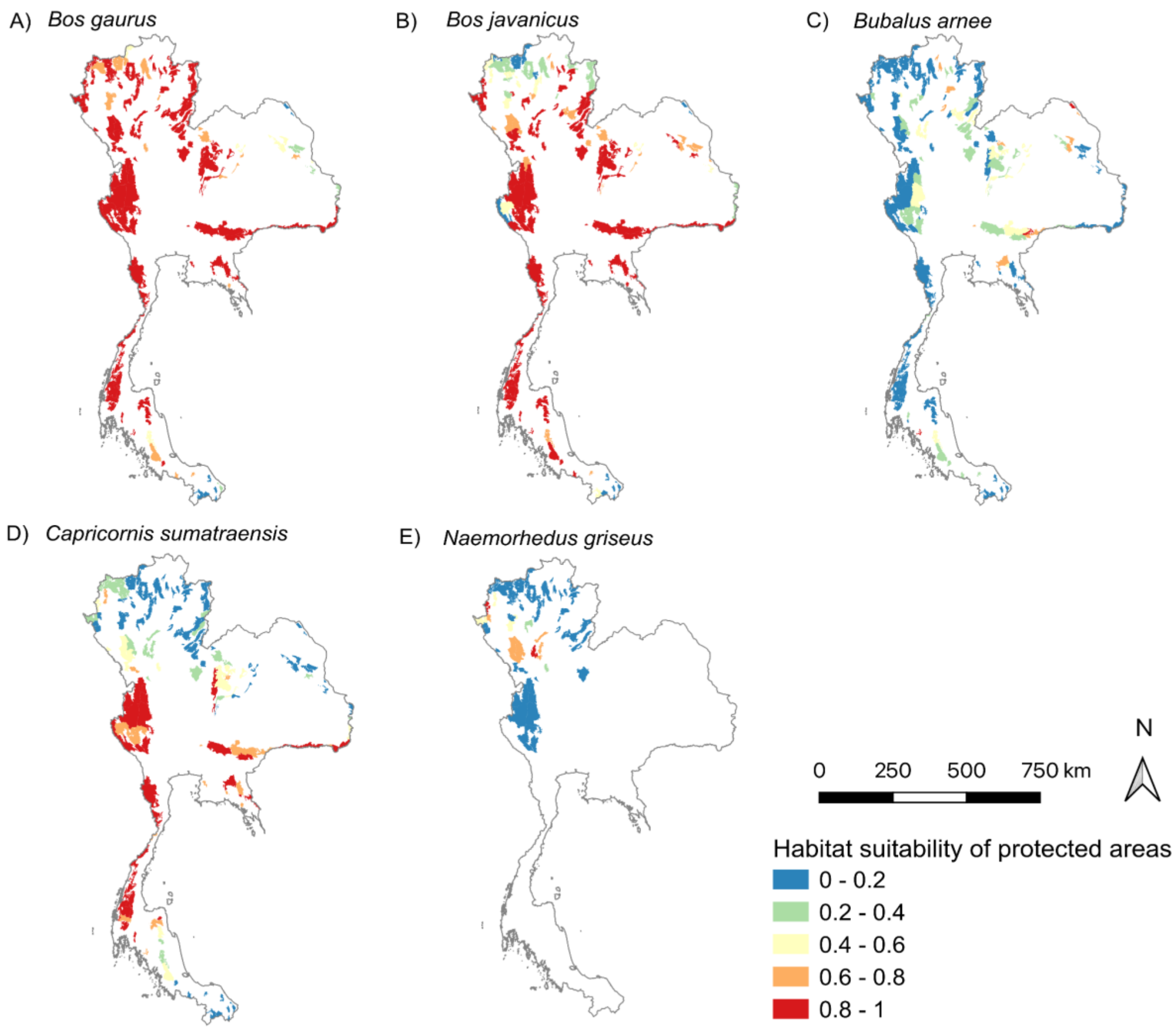


Figure S10 The proportion of suitable areas range from 0 (unsuitable) to 1 (high proportion of suitable) focusing on Thailand PAs for five species and their best performing models: (A) Guar (*Bos gaurus*), (B) Banteng (*Bos javanicus*), (C) wild water buffalo (*Bubalus arnee*), (D) mainland serow (*Capricornis sumatraensis*), and (E) Chinese goral (*Naemorhedus griseus*).
